# The fate of deleterious variants in a barley genomic prediction population

**DOI:** 10.1101/442020

**Authors:** TJY Kono, C Liu, EE Vonderharr, D Koenig, JC Fay, KP Smith, PL Morrell

## Abstract

Targeted identification and purging of deleterious genetic variants has been proposed as a novel approach to animal and plant breeding. This strategy is motivated, in part, by the observation that demographic events and strong selection associated with cultivated species pose a “cost of domestication.” This includes an increase in the proportion of genetic variants where a mutation is likely to reduce fitness. Recent advances in DNA resequencing and sequence constraint-based approaches to predict the functional impact of a mutation permit the identification of putatively deleterious SNPs (dSNPs) on a genome-wide scale. Using exome capture resequencing of 21 barley 6-row spring breeding lines, we identify 3,855 dSNPs among 497,754 total SNPs. In order to polarize SNPs as ancestral versus derived, we generated whole genome resequencing data of *Hordeum murinum* ssp. *glaucum* as a phylogenetic outgroup. The dSNPs occur at higher density in portions of the genome with a higher recombination rate than in pericentromeric regions with lower recombination rate and gene density. Using 5,215 progeny from a genomic prediction experiment, we examine the fate of dSNPs over three breeding cycles. Average derived allele frequency is lower for dSNPs than any other class of variants. Adjusting for initial frequency, derived alleles at dSNPs reduce in frequency or are lost more often than other classes of SNPs. The highest yielding lines in the experiment, as chosen by standard genomic prediction approaches, carry fewer homozygous dSNPs than randomly sampled lines from the same progeny cycle. In the final cycle of the experiment, progeny selected by genomic prediction have a mean of 5.6% fewer homozygous dSNPs relative to randomly chosen progeny from the same cycle.

**Author Summary:** The nature of genetic variants underlying complex trait variation has been the source of debate in evolutionary biology. Here, we provide evidence that agronomically important phenotypes are influenced by rare, putatively deleterious variants. We use exome capture resequencing and a hypothesis-based test for codon conservation to predict deleterious SNPs (dSNPS) in the parents of a multi-parent barley breeding population. We also generated whole-genome resequencing data of *Hordeum murinum*, a phylogenetic outgroup to barley, to polarize dSNPs by ancestral versus derived state. dSNPs occur disproportionately in the gene-rich chromosome arms, rather than in the recombination-poor pericentromeric regions. They also decrease in frequency more often than other variants at the same initial frequency during recurrent selection for grain yield and disease resistance. Finally, we identify a region on chromosome 4H that strongly associated with agronomic phenotypes in which dSNPs appear to be hitchhiking with favorable variants. Our results show that targeted identification and removal of dSNPs from breeding programs is a viable strategy for crop improvement, and that standard genomic prediction approaches may already contain some information about unobserved segregating dSNPs.

## Introduction

Gains from selection in plant and animal breeding could be improved through a better understanding of the genetic architecture of complex traits. One current source of debate is the relative frequency of genetic variants that contribute to complex traits. At mutation-drift equilibrium, the majority of genetic variants segregating in a population are expected to be rare [1,2]. If a genetic variant affects a phenotype, it is more likely to be subject to selection, with the strength of selection proportional to the magnitude of phenotypic impact [3]. Since most new mutations with a phenotypic impact are expected to be deleterious [4–6], variants contributing to complex trait variation will likely be under purifying selection [7,8]. Thus, a substantial portion of genetic variants that affect phenotypes may occur as “rare alleles of large effect” (RALE) [3]. Consistent with the RALE hypothesis, association mapping studies find evidence that rare alleles have larger estimated phenotypic effects than common alleles [9]. Because of their frequency, rare alleles are more difficult to associate with a phenotype. Alleles with relatively large effects on phenotype are more readily detected [10,11] but are unlikely to be representative of the majority of genetic variants that contribute to phenotypic variation [10,12].

Segregating variants that affect fitness are more likely to be deleterious than beneficial [13] and are thus more likely to be under purifying selection. Consistent with this postulate, low frequency genetic variants in human populations are enriched for amino acid replacements [e.g., 14], which likely have direct effects on protein function. The effect of individual dSNPs on fitness is expected to be small, but in aggregate their impact may be substantial [cf. 13]. Domesticated plants and animal populations have often experienced reductions in effective population size and strong selection associated with domestication and improvement that could result in exacerbated effects of deleterious variants as a genetic “cost of domestication” [15]. Empirical evidence from a variety of organisms appears to support this conjecture, with comparisons in cassava [16], dogs [17], grapes [18], and rice [19] showing evidence of an increased proportion of both fixed and segregating dSNPs relative to wild progenitors [see also 20,21].

Putative dSNPs can be readily identified based on phylogenetic conservation, particularly for coding polymorphisms [22,23]. SNPs that are phenotype-changing in *Arabidopsis thaliana* are more likely to annotate as deleterious than “tolerated” (less conserved) amino acid changing SNPs at similar frequencies [24]. Indeed, a number of putatively causative amino acid changing SNPs that contribute to agronomic phenotypes annotate as deleterious [25]. However, individual inbred lines for many cultivated species carry hundreds to thousands of dSNPs [25,26]. The vast majority of dSNPs occur at low frequency [19,25] and thus are unlikely to serve as the primary causative variants for essential agronomic traits. Because of their relative ease of identification, elimination of dSNPs either through selection against them in aggregate [20,27–29] or through targeted replacement of individual dSNPs [27,28,30] provides a potential means of crop improvement.

The phenotypic consequences of dSNPs is determined by their relative degree of dominance, the proportion of variants that occur in the homozygous state, and the fitness effects of individual dSNPs [17,20,31–34]. Additionally, the genomic locations of dSNPs is an important factor in how effective purifying selection can be in culling them from populations. This is due to recombination rate variation placing limits on the efficacy of purifying selection [35,36]. A larger proportion of variants may be deleterious in low recombination regions of a genome [13] as has been observed in sunflower [26], rice [19], and soybean [25]. There is evidence from studies of humans that variants in low recombination regions may have larger fitness consequences or explain more of the variation for quantitative traits [37,38].

Modern breeding programs use genome-wide prediction approaches which are designed to integrate large numbers of markers in the estimation of phenotypic values for quantitative traits [39]. This typically involves the use of a training panel of individuals with both genotypic and phenotypic information. Prediction and selection can be performed in a panel of related individuals with only genotypic data. There is evidence that the probable effect of genetic variants on quantitative phenotypic variation can vary by functional class and that prediction accuracy can be improved through differential weighting of variants [34,40].

The purpose of this study is to assess the fate of dSNPs in a breeding population subject to genomic prediction and selection. The experimental barley breeding population was developed at the University of Minnesota [41]. Genomic prediction was used to select lines with improvements in yield and resistance to the fungal disease Fusarium head blight (FHB), two unfavorably correlated quantitative traits. Phenotypic data was collected for yield, deoxynivalenol (DON) concentration (a measure of severity of fungal infection), and for plant height, which was not under selection. The population showed gains in both yield and FHB resistance over three cycles of crossing and selection, with an index of yield and reduced DON concentration showing consistent gain over cycles [41]. The pedigreed design brings the rarest variants to ∼3% frequency, thus improving the potential to assess the contributions of putative dSNPs to agronomic phenotypes. The major questions we seek to address are: (1) How common are putative dSNPs in elite barley breeding material? (2) Are putative dSNPs uniformly distributed across the genome or concentrated in genomic regions with lower rates of recombination?; and (3) What is their fate through rounds of selection and breeding gain in an experimental breeding population? We also make use of a linear mixed model to estimate the proportion of phenotypic variance that can be explained based on SNPs genotyped in our panel or imputed from parents onto progeny. We find a genomic region associated with agronomic traits in which dSNPs may be hitchhiking due to strong selection in this population.

## Results

### Summary of Resequencing Data

We make use of exome capture resequencing to identify nucleotide sequence variants in 21 barley breeding lines from three barley breeding programs (S1 Table). The 5,215 progeny in the experiment were genotyped using a 384 SNP Illumina assay [41]. Based on observed genotypes in progeny in the known pedigree, we track the fate of genotyped and imputed SNPs through three breeding cycles (S1 Fig). All lines are part of a genomic prediction experiment [41] where sets of progeny were selected based on genomic prediction for yield and fungal disease resistance. A second pool of progeny was drawn at random in each cycle and subject to the same field testing for yield and disease resistance as selected progeny.

The 21 parents (Cycle 0) in the experiment were subjected to exome capture resequencing, resulting in the identification of 497,754 SNPs. Of these, 407,285 map to portions of the reference genome that could be assigned to barley chromosomes and are subject to further analysis (Table 1). The intersections of three deleterious annotation approaches identified 3,855 dSNPs at 62,826 nonsynonymous sites, including 1,877 early stop codons in the founding parents. More of the the dSNPs are private to North Dakota lines than to the other programs (Table 2), which has more private SNPs across classes. The numbers of dSNPs is remarkably similar among lines with a mean of 677.67 (± 16.51), though the number of dSNPs private to a line varies more dramatically (from 11 to 172) (S2 Table). The unfolded site frequency spectrum (SFS) for 283,021 SNPs with inferred ancestral state indicates that dSNPs in the founders occur primary in the rarest frequency classes (Figure 1), a trend that is also evident among all variants in the folded SFS (S2 Fig).

**Table 1.**
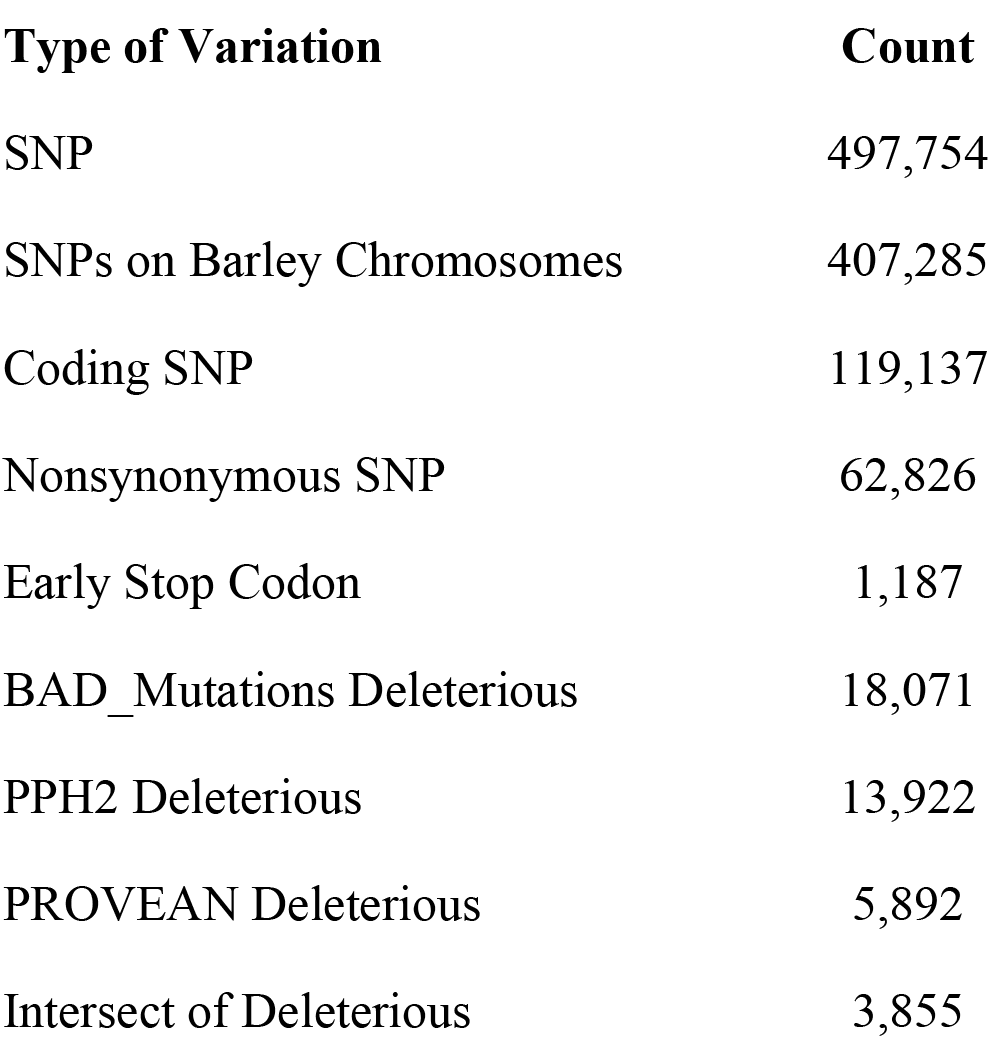
Summary of SNPs identified in exome capture resequencing of parental accessions.

| Type of Variation | Count |
| --- | --- |
| SNP | 497,754 |
| SNPs on Barley Chromosomes | 407,285 |
| Coding SNP | 119,137 |
| Nonsynonymous SNP | 62,826 |
| Early Stop Codon | 1,187 |
| BAD_Mutations Deleterious | 18,071 |
| PPH2 Deleterious | 13,922 |
| PROVEAN Deleterious | 5,892 |
| Intersect of Deleterious | 3,855 |

**Table 2.** The number of private SNPs per population across three classes of genic variants.

| Breeding Program | Synonymous | Nonsynonymous | Deleterious |
| --- | --- | --- | --- |
| Busch Ag | 3,578 | 3,906 | 271 |
| Minnesota | 7,930 | 9,031 | 500 |
| North Dakota | 13,581 | 15,166 | 1,019 |

**Figure 1.**
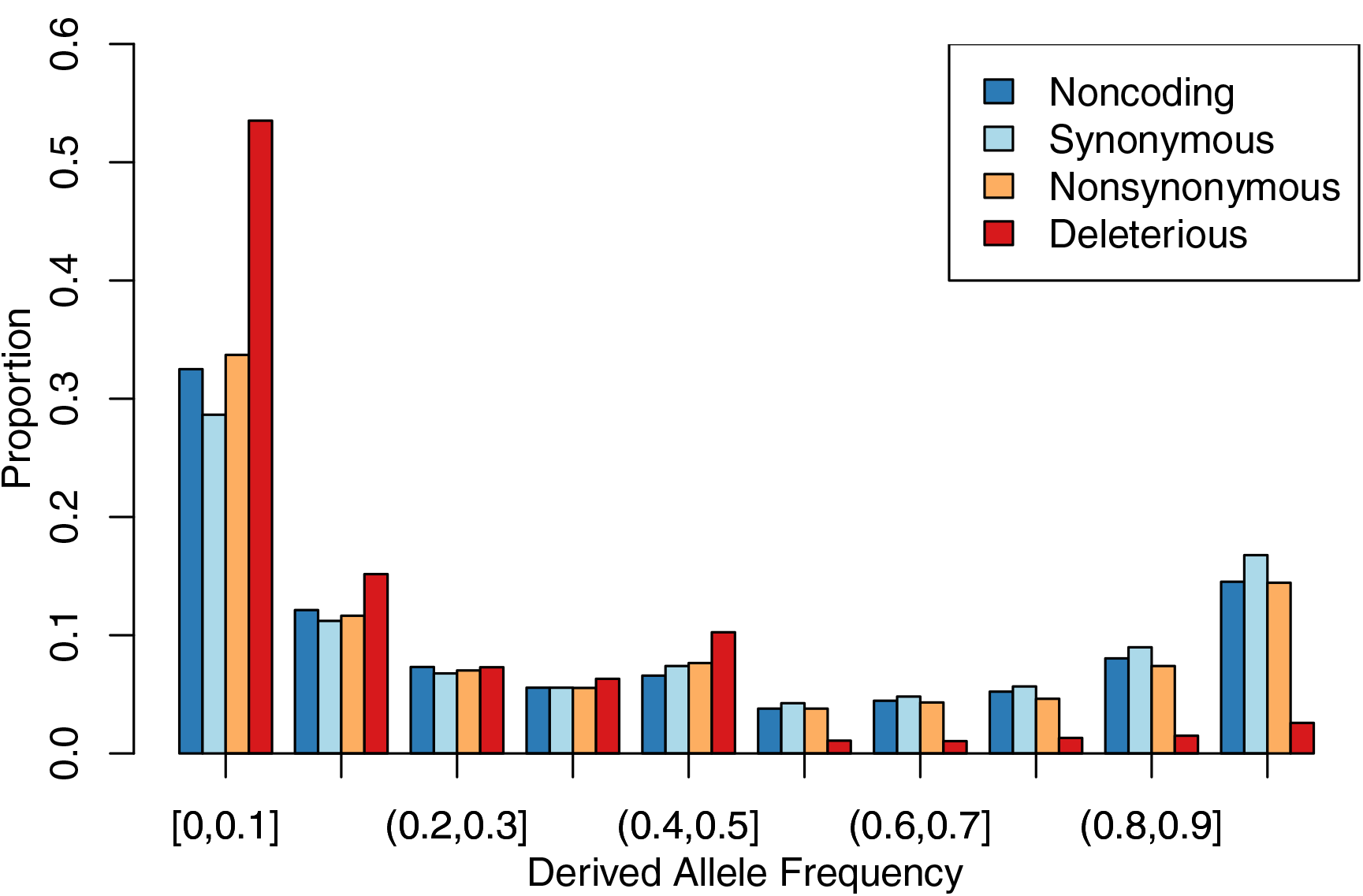
Derived site frequency spectra, for 283,021 SNPs in the parental founders of the genomic prediction population. Ancestral state was based on majority state from *Hordeum murinum spp. glaucum* resequencing mapped to the Morex assembly. (For all SNPs, including those with no inferred ancestral state see Figure S2.) “Noncoding” refers to SNPs in regions that do not code for proteins, “Synonymous” refers to SNPs in coding regions that do not alter an amino acid sequence, and “Nonsynonymous” refers to SNPs that alter the amino acid sequence. SNPs listed as “Deleterious” are the intersect of variants that annotate as deleterious in each of three approaches.

SNP density was highest along chromosome arms and lower in pericentromeric regions (S3 Fig), consistent with the reports of the distribution of gene density [42,43]. Using pericentromeric regions as defined based on barley recombination rate and gene density reported by [42], we identify 71,939,192 bp (81.3%) of capture targets in euchromatic regions and 16,511,574 bp (18.7%) in pericentomeres (A BED file of positions covered by exome capture is available at http://conservancy.umn.edu/XXXX). Codon density was similar within exome capture from the two regions. Euchromatic regions include 6,945,584 bp (81.3%) of codons within capture targets and 1,592,281 bp (18.7%) in pericentromeric regions. The euchromatic regions include 401,148 (86.6%) of SNPs versus 62,060 (13.4%) of SNPs in pericentromeres. This resulted in 3,331 (87.7%) dSNPs in euchromatin and 466 (12.3%) dSNPs in pericentromeric regions. Thus the proportion of dSNPs per codon is lower in the pericentromere than in higher recombination regions (Figure 2).

**Figure 2.**
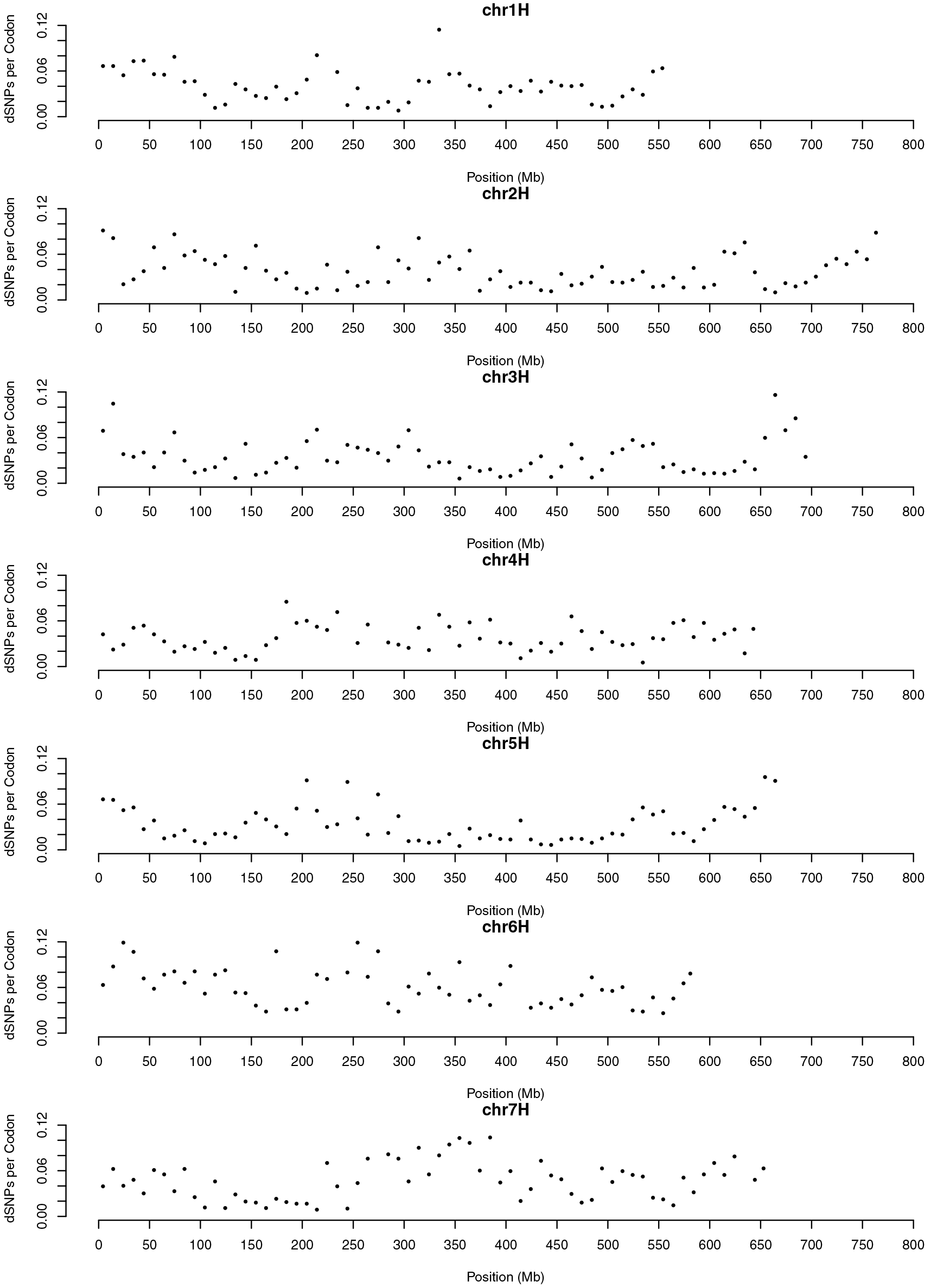
The number of dSNPs per covered codon in 1Mb windows across the barley genome. The light grey shading shows the pericentromeric region, and the dark grey shading show the centromere.

To infer the ancestral state of variants in cultivated barley, we performed whole genome resequencing of *H. murinum* ssp. *glaucum*, yielding 371,255,479 reads. A divergence rate setting of 3% in Stampy [44] resulted in the largest percentages of reads mapping to the reference genome. Genome-wide coverage was estimated as 37 X. This permitted estimation of ancestral state for 283,021 or 69.5% of barley SNPs. Results of ancestral state inference by functional class of variants is show in S3 Table.

### Genotyping Data

The final dataset used for analysis consisted of 5,215 individuals. Of the 384 SNPs on the custom Illumina Veracode assay [41], four were eliminated because of errors in Mendelian inheritance between parents and progeny. Three SNPs with >20% missing genotypes were also excluded, resulting in 377 SNPs segregating among progeny (S3 Fig). For 16 SNPs, either genetic or physical positions needed to be interpolated from flanking SNPs (see Supplemental Text). The parental lines and progeny produced an average of 366.5 (± 40.1) genotyped SNPs. Pairwise diversity averaged 0.32 across cycles, with observed heterozygosity between 8 and 15% in C1 through C3 (S4 Table).

Using the 377 genotyped Veracode SNPs, we imputed genotypes for all variants in the pedigreed populations using the program AlphaPeel [45]. Imputed genotypes are reported in AlphaPeel output as the expected dosage of the non-reference allele at each site. Recombination probabilities are modeled from interpolated genetic distances between observed markers with known genetic distances [45]. Both the unfolded (S4 Fig) and folded SFS (with all variants) (S5 Fig), demonstrate that dSNPs remain at low frequency across generations in the population. Average pairwise diversity for SNPs resequenced in the founder lines and imputed onto progeny was ∼0.19 for synonymous SNPs and ∼0.12 for dSNPS, with noncoding and nonsynonymous having intermediate levels of diversity (S5 Table).

### Putatively Deleterious SNPs and Phenotypic Variation

A total of 676 of the 5,215 individuals have phenotypic data for grain yield, DON concentration, and plant height. Yield increased and average DON concentration decreased over three cycles of selection (Figure 3). An index of yield and DON concentration showed steady improvement in each cycle [41]. Plant height, which was not subject to selection in this population, increased over the course of the experiment (Figure 3). The number of putative dSNPs that were homozygous for the derived allele within an individual is significantly correlated with all three measured phenotypes (Figure 4). Yield is negatively correlated with the number of homozygous derived SNPs across all classes. The correlation is greatest for noncoding (the largest class of) SNPs. Based on a product moment correlation, the correlation is significant at *p* < 0.05 for noncoding and nonsynoymous, and at *p* < 0.001 for dSNPs (Table 3). For DON concentration and plant height, where larger values are the less desirable trait, the correlations with the number of homozygous derived SNPs are positive. These correlations are significant with the notable exception of DON and dSNPs (Table 3).

**Figure 3.**
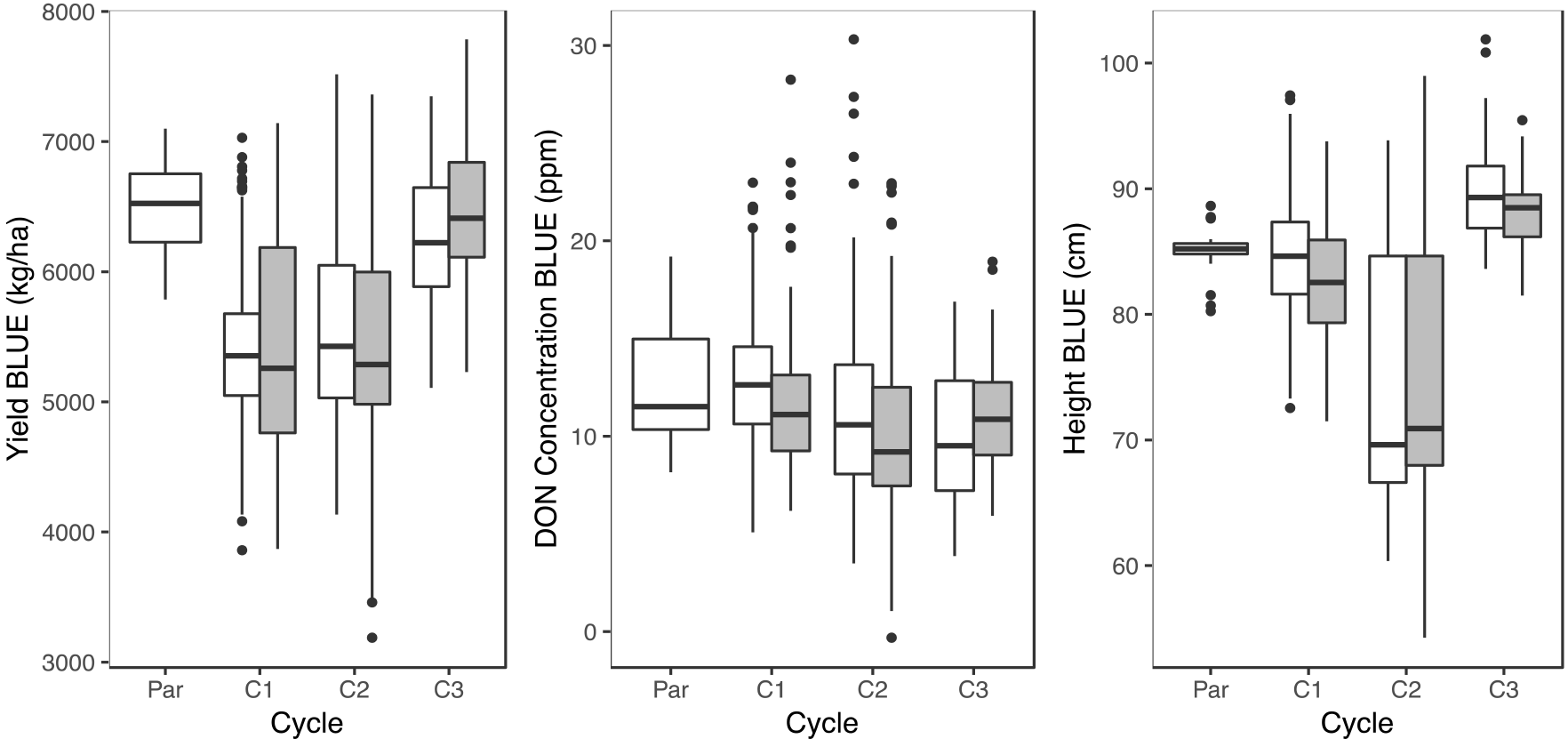
Plots of yield (A) and DON concentration (B) and plant height (C) data collected on the experimental population. Values for check lines, founding parents (Cycle 0), and each of three cycles are shown (C1 - C3). In C1 - C3, randomly selected lines are shown in white, and lines selected based on genomic estimated breeding dvalues are shown in grey. Data shown is the linear unbiased estimates (BLUEs) for individual lines based on yield, DON, and plant height observations at five year-locations.

**Figure 4.**
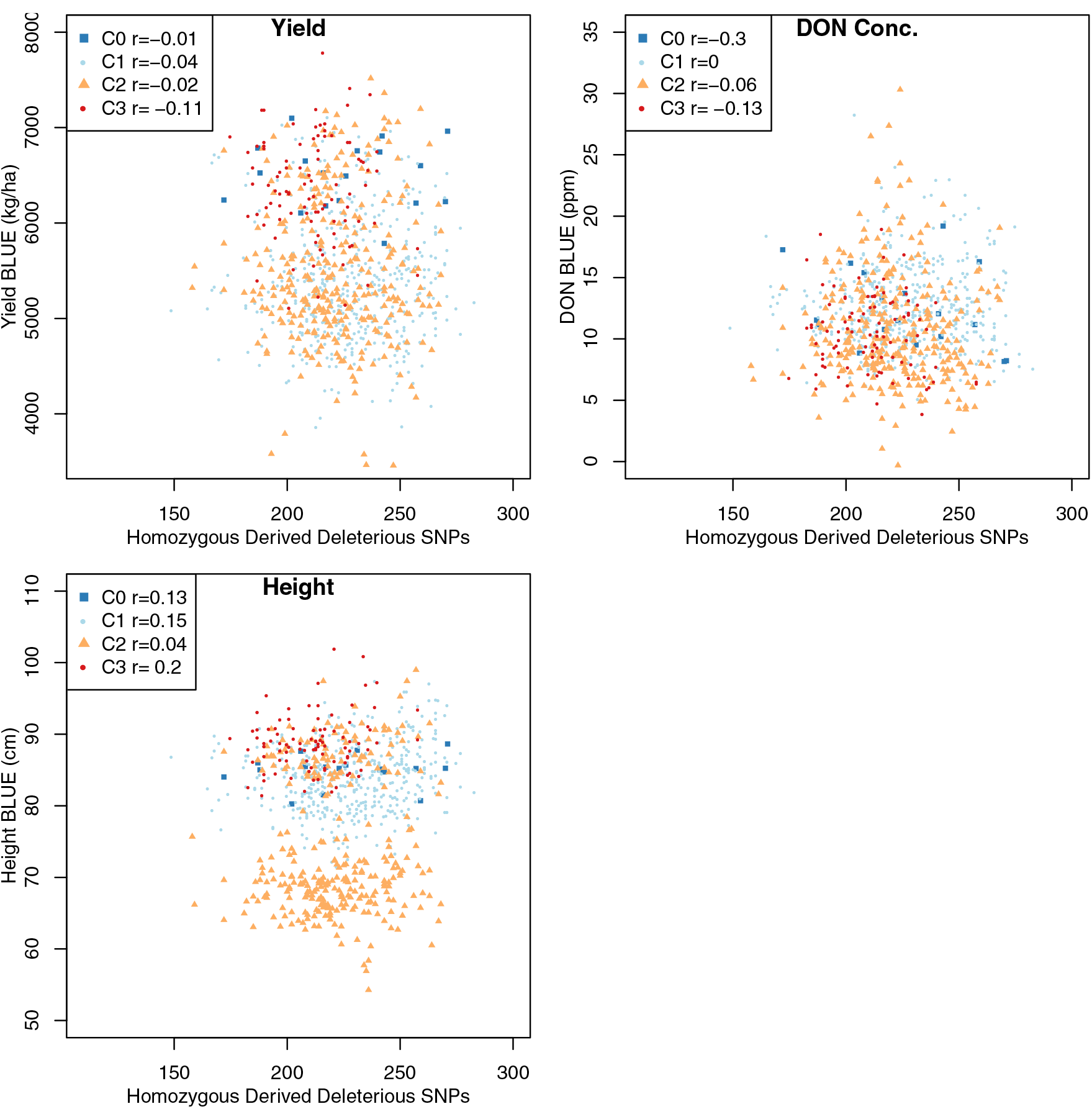
The number of homozygous dSNPs in each cycle of the experiment compared to the BLUE for yield, DON concentration, and plant height. Values are shown for Cycle 0 to Cycle 3.

**Table 3.**
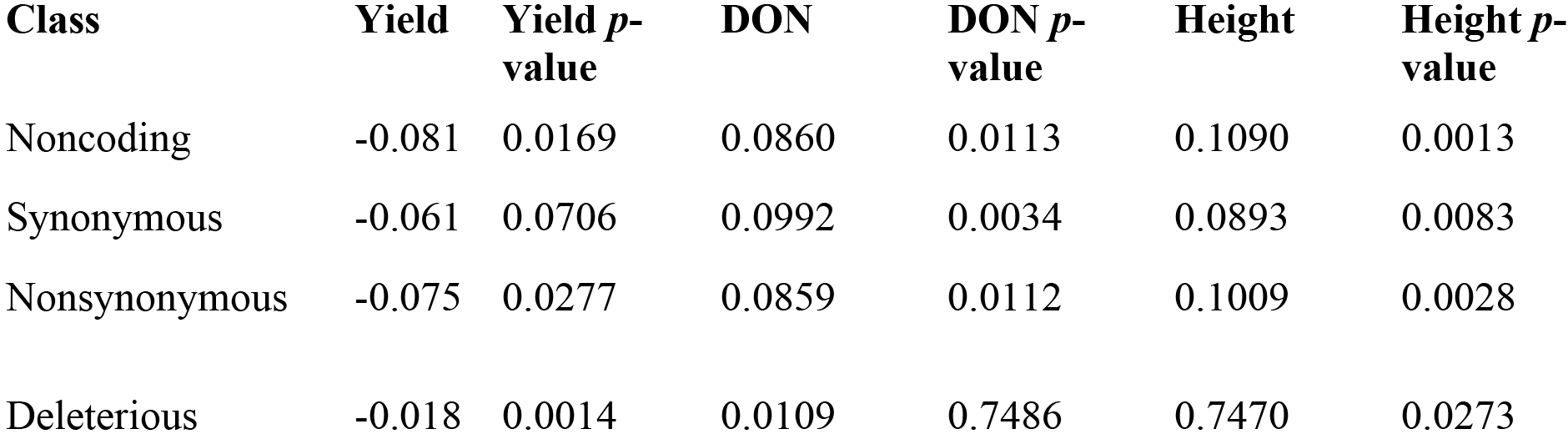
The correlation between the number of homozygous derived genotypes and the phenotypes across each functional class of variants.

The proportion of phenotypic variance explained by all genotypes jointly, also referred to as “SNP heritability” was estimated using a linear mixed model implemented in GEMMA [46]. The removal of SNPs with a minimum minor allele frequency (MAF) of ≤ 1% resulted in the inclusion of 357 of the 377 SNPs genotyped in all progeny. Heritability estimates for this SNP set were 0.198 for yield, 0.357 for DON concentration, and 0.237 for height.

Among the SNPs directly genotyped in the progeny, three (11_10196, 11_20422, and 11_20777) were identified as contributing to yield with a *p* < 0.01. The first SNP had a favorable effect on yield while the latter two SNPs were associated with reduced yield and with the favorable trait of reduced DON concentration. All three SNPs are at relative high minor allele frequencies (∼ 0.3 - 0.4) and increase in frequency from C0 to C3. All three occur in chromosomal regions on 2H and 4H previously identified as under selection in Minnesota barley breeding lines subject to introgression for increased Fusarium head blight resistance [47]. A region on chromosome 4H (18.7 - 35.8 Mb) contributes six of eight associations with *p* < 0.01 for DON concentration and overlaps with a region of the genome that [47] demonstrated had been subject to strong selection for Fusarium resistance. The region covers ∼ 2.6% of the 647 Mb of chromosome 4H and includes 110 annotated genes. Fifteen dSNPs were identified in this interval. For eight dSNPs with unambiguous ancestral state, frequencies were maintained or increased over breeding cycles, resulting in a mean DAF of 0.60 in C3. The dSNPs were included in a major haplotype contributed by one of three founders, FEG153-58, FEG154-47, or FEG175-57, all from the Minnesota breeding program.

For linear mixed model analysis using SNPs identified in exome capture, the ≥ 1% frequency threshold resulted in retention of 419,956 SNPs (86% of all SNPs). Heritability estimates were 0.250 for yield, 0.514 for DON concentration, and 0.358 for plant height. These values are consistent with previous estimates: a study of a two-row barley double haploid population grown across 25 locations reported average yield heritability of 0.35 and plant height of 0.33 [48]. Heritability for DON accumulation has been estimated as 0.46 in a separate study of crosses between two-row and six-row barley [49].

### Change in SNP Frequency over Cycles

Using the parental assignment of genomic segments in the progeny, it is possible to track changes in frequency for segregating variation across various functional classes of SNPs. While all classes of SNPs became more homozygous over generations, dSNPs are lost from the population more frequently than synonymous SNPs (Table 4). Out of 37,766 synonymous SNPs with unambiguous ancestral state (required for dSNPs to infer which variant is likely deleterious) identified in the parents, 30,481 (80.7%) were still segregating in Cycle 3. Of the 1,913 dSNPs identified with unambiguous ancestral state, 1,278 (66.8%) were segregating in Cycle 3. However, this measure does not account for lower average derived allele frequencies for dSNPs. If measured as relative fold change in derived allele frequency, dSNPs are more frequently decreasing in frequency (Figure 5). The median change in DAF is −0.25 for dSNPs and closer to zero for all other classes (Table 5). Slightly more than half of variants showed decreased DAF over breeding cycles, but this trend is observed at 0.627 of dSNPs. When using the pedigree to establish expectations for the allele frequencies in each cycle, we still observe a preferential loss of dSNPs as segregating variation (S4 Fig; Table 4). When considering the variants with an inferred ancestral state, dSNPs have a larger proportion of variants that fix for the ancestral state than other classes of variants (S6 Fig). Fold change across the genome for individual classes of variants can be seen in S7 Fig). Of the 1,913 dSNPs with inferred ancestral state, 621 (32.5%) are fixed for the ancestral allele, while 14.6%, 2.8%, and 2.7% of noncoding, synonymous, and nonsynonymous SNPs were fixed for the ancestral allele, respectively.

**Table 4.** The proportion of variants in the founding parents from each functional class that were lost, segregating, or fixed in progeny in Cycle 03.

| **Class** | **Segregating** | **Lost** | **Fixed** |
| --- | --- | --- | --- |
| Noncoding | 120,289 | 22,083 | 6,774 |
| Synonymous | 30,481 | 5,282 | 2,003 |
| Nonsynonymous | 25,633 | 6,047 | 1,610 |
| Deleterious | 1278 | 621 | 14 |

**Figure 5.**
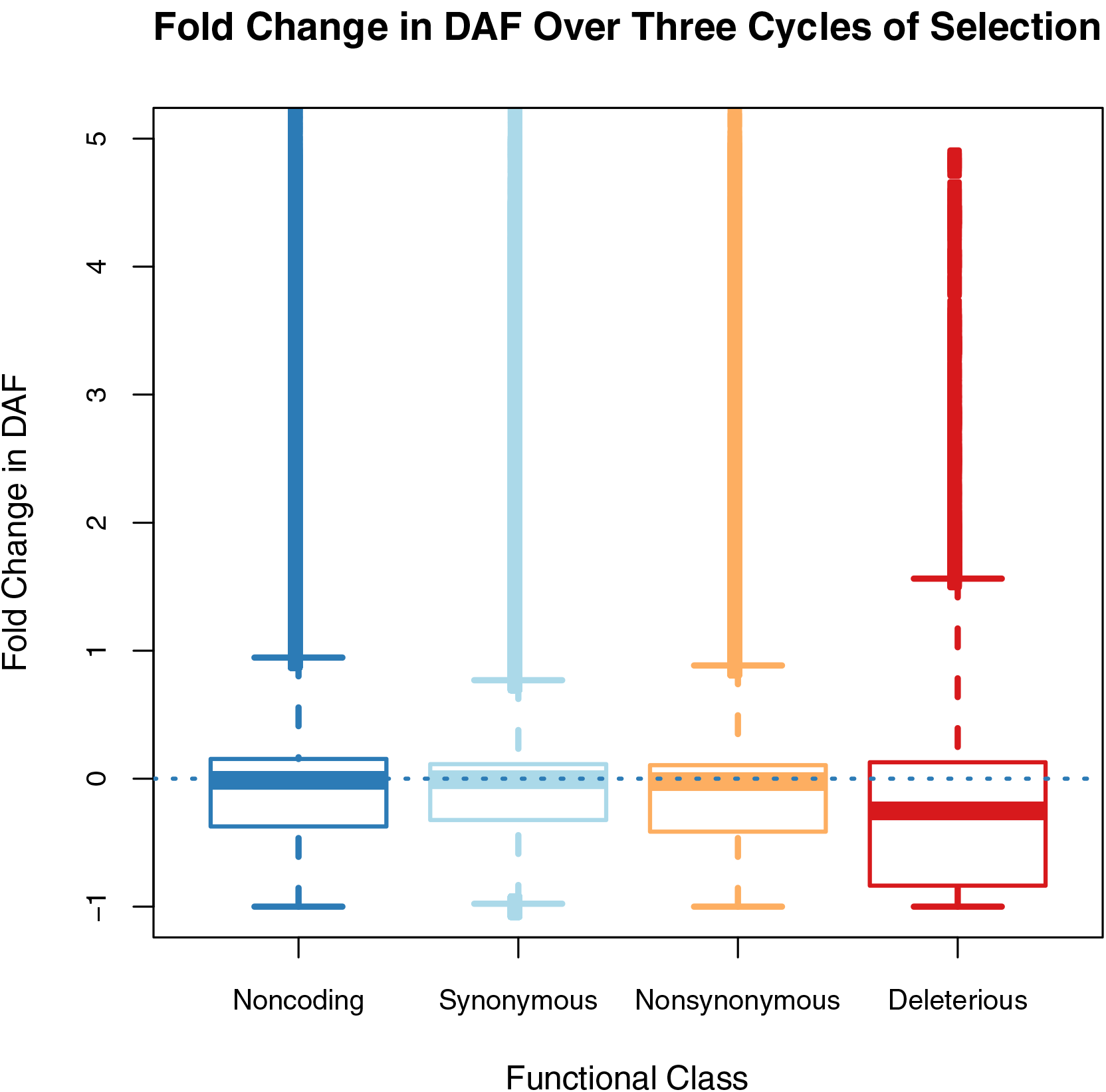
The fold change in derived allele frequency (DAF) in SNPs from four classes of variants. The nonsynonymous class includes only SNPs determined to be ‘tolerated’ based on deleterious variant annotation.

**Table 5.** The change in SNP frequency by class. Nonsynonymous includes SNPs that are amino acid-changing but are annotated as “tolerated.” The values reported are the median change in derived allele frequency (DAF), and the proportions of SNPs increasing or decreasing over the three breeding cycles.

| **Class** | **Median Change in DAF** | **Increasing** | **Decreasing** |
| --- | --- | --- | --- |
| Noncoding | -0.0125 | 0.460 | 0.527 |
| Synonymous | -0.0075 | 0.465 | 0.519 |
| Nonsynonymous | -0.0220 | 0.439 | 0.543 |
| Deleterious | -0.2510 | 0.343 | 0.627 |

The number of homozygous derived dSNPs is reduced in each cycle, but is reduced more dramatically for the lines selected for yield and reduced DON concentration than for random chosen lines from the same cycle (Figure 6). In other classes of variants, selected lines tend to have slightly more homozygous derived variants than random chosen lines; across classes of SNPs, derived homozygous variants become less frequent over cycles.

**Figure 6.**
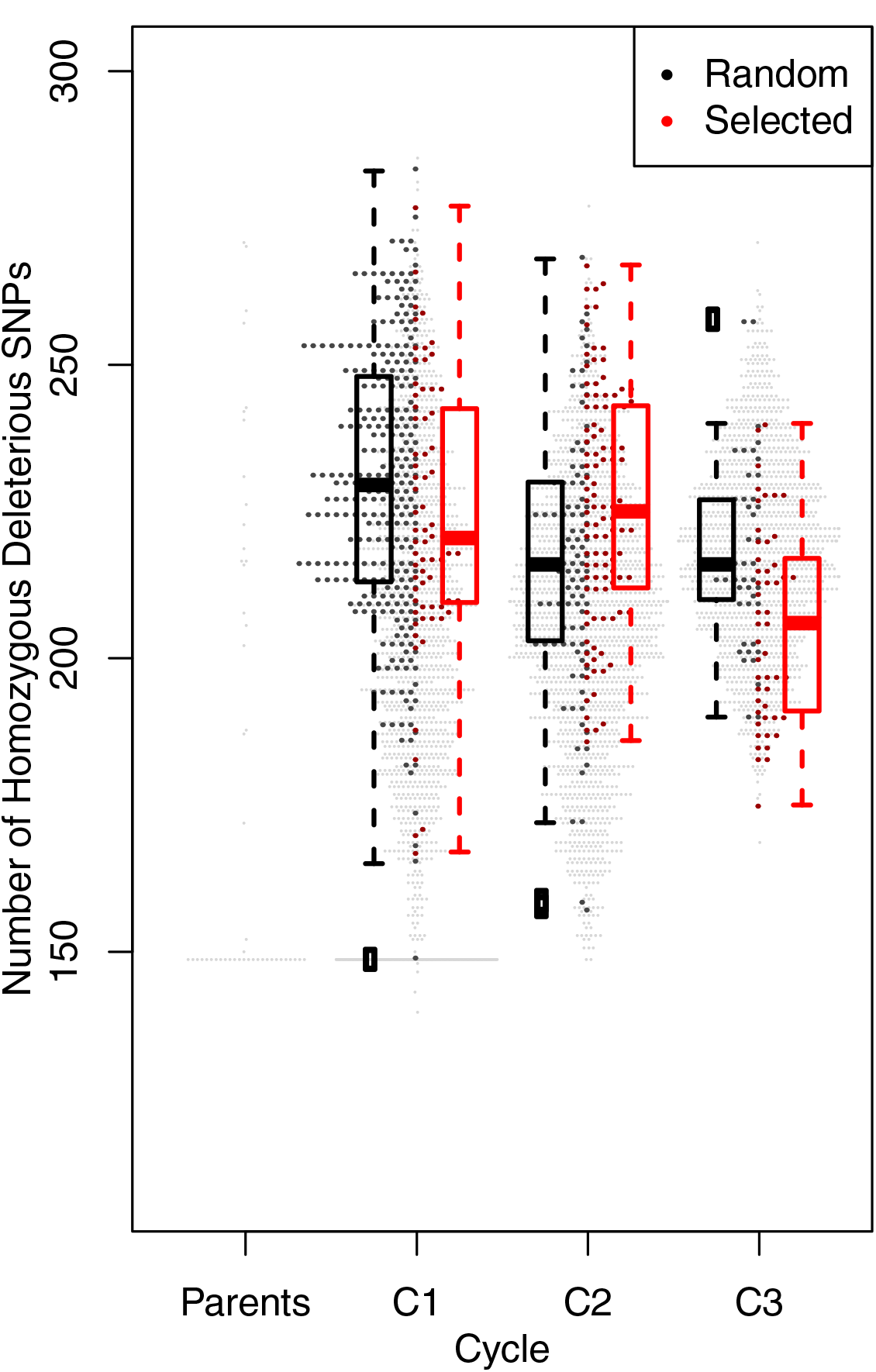
Number of dSNPs in the homozygous state in parents and progeny over three breeding cycles, C1 - C3. Values for all individuals are shown, with random samples in C1 - C3 in black and selected samples in red. Boxplots summarize the data for each partition of samples.

With regard to homozygous derived dSNPs, the difference in selected and random lines differed by cycle. For Cycle 1, selected lines had a mean of 224.25 (± 22.72) homozygous dSNPs relative to 229.05 (± 22.82) in randomly chosen lines, a difference that was not significant in a one-sided t-test, *p* = 0.060. The dSNP mean homozygosity was a slight decrease from 225.50 (± 28.87) homozygous dSNPs in founders in Cycle 0. Selection in Cycle 2 saw dramatic reduction in DON concentration but little change in yield (Figure 3), [see also41]. In that generation, selected lines averaged more dSNPs than randomly chosen lines, 225.73 (± 19.87) versus 216.34 (± 2139). Cycle 3 progeny showed yield improvement, with minimal change in DON. The difference in selected and random chosen lines for mean dSNPs was large, with 205.86 (± 16.43) versus 218.02 (± 15.52), with *p* = 0.00017 in a one-sided t-test. The number of homozygous dSNPs over generations changes more dramatically than the dosage of dSNPs in individual lines (S8 Fig), consistent with effects of dSNPs being primarily recessive.

## Discussion

We examined the fate of multiple classes of variants in a population subjected to genomic prediction and selection for two unfavorably correlated quantitative traits over three cycles. Selection was based on genomic prediction from a genome-wide set of 384 SNPs genotyped in all progeny. This selection did not make use of any information on functional annotation of variants. We identify 3,855 putative dSNPs segregating in protein coding regions; most of these SNPs are at low frequency in the founding parents (Figure 1; S2 Fig) and on average, decrease slightly in frequency over the course of the experiment (S4 Fig, S5 Fig). The highest yielding progeny in the population carry fewer dSNPs than progeny drawn at random (Figure 6).

Over three cycles of intercrossing and selection, the proportion of dSNPs occurring in the highest derived frequency class (S4 Fig) or reaching fixation (Table 4) is notably lower than other classes of SNPs. Taken together, these lines of evidence suggest that dSNPs that contribute to a diminution of yield are selected against despite the limitations of population size and the countervailing effects of selection on predicted yield and FHB resistance.

Though progeny were selected for both predicted yield and FHB resistance, lines selected based on genomic breeding value typically have a lower total dosage of dSNPs (including SNPs in both the heterozygous and homozygous state) (S8 Fig), and fewer dSNPs in the homozygous state (Figure 6). The reduction in homozygous variants per line is consistent with the majority of dSNPs constituting recessive, loss of function changes. The reduction in the number of homozygous dSNPs occurs over successive generations in the experiment, resulting in a significant negative correlation between both yield and the number of homozygous SNPS, including dSNPs. A larger number of homozygous derived SNPs is associated with higher DON, the undesirable state. The correlation of DON concentration and dSNPs is not statistically significant (Table 3). This is consistent with the expectation that dSNPs are more likely to be predictive to fitness-related phenotypes such as yield [16,30,34]. Plant height was not under selection, but increasing plant height is generally not desirable. It was positively correlated with the number of homozgyous derived SNPs (Table 3).

The barley genome includes large pericentromeric regions with minimal crossover [42,43] (S3 Figure). Based on our exome capture resequencing, these regions harbor fewer dSNPs per codon than the distal arms of chromosomes (Figure 2). This should not be taken as evidence that linked selection in these regions is unimportant, but rather that gene density plays an important role in determining the distribution of dSNPs within coding regions. Previous studies have suggested dSNPs occur at a higher frequency in lower recombination regions of the genome in sunflower [26], rice [15,19], and soybean [25]. Evidence for this phenomenon in maize is mixed, with no evidence for higher mutational load reported by [31] whereas it was identified by [50]. Comparison among studies is made more difficult by differences in approaches for dSNP annotation and the sequence diversity statistics used as a point of comparison (e.g., density of synonymous SNPs) [see 20,19]. There may also be a weaker relationship between recombination and diversity in predominantly self-fertilizing species [51]. An implication is that for barley, and perhaps other species, the majority of dSNPs occur in genomic regions where crossover rates are relatively high. Thus many dSNPs can potentially be removed from populations based on the action of crossover and independent assortment.

This study involved simultaneous prediction and selection on two quantitative traits that are unfavorably correlated. This represents a somewhat realistic scenario for many applications of genomic prediction. Based on linear mixed model analysis of marker-trait association, we identified a 17.1 Mb region on chromosome 4H that contributed to reduced disease severity but also had negative impacts on yield. This region had been previously identified in a selection mapping study for FHB resistance [47]. Selection for variants in this region contributed to improved disease resistance but also provided the opportunity for at least 15 identified dSNPs to be maintained in the population. For eight of those variants where the derived (and likely deleterious state) is unambiguous, there is evidence of hitchhiking to higher frequencies.

The identification and weighting of deleterious variants in a genomic prediction framework appears to be promising path for improving phenotypic prediction [27,34]. While we observed little difference in the number of dSNPs per line, the number of private dSNPs varied dramatically, providing the opportunity to select progeny with fewer rare and potentially deleterious variant than either parent. It should be noted that the fitness effects of individual deleterious variants in crops remains largely unknown and indeed, the shape of the distribution of fitness effects of all variants is a challenging quantity to estimate [52,53]. The proportion of variants with large effects on fitness could impact genomic prediction strategies. As with any examination of a complex trait, sample sizes for phenotyped individuals are likely to limit power to detect effects among classes of variants. Also, because deleterious variants are completely commingled with other classes of variants, limits on recombination within a population limit the degree to which the effects of deleterious variants can be isolated. Given these caveats, the potential to readily identify a class of fitness-related variants that can be subject to selection holds considerable promise for phenotypic prediction.

## Materials and Methods

### Population Design

Our experimental population consists of spring, six-row, malting barley adapted to the Upper Midwest of the United States. Three breeding programs (Busch Agricultural Resources, Inc., North Dakota State University, and University of Minnesota) contributed the 21 founders of the population, denoted as Cycle 0 (C0) (S1 Table; S1 Figure). Founders were used to produce 45 crosses (pedigrees available at http://conservancy.umn.edu/XXXX). F_1_ progeny from each of the crosses were self-fertilized to the F_3_ generation, resulting in 1,080 F_3_ progeny, denoted as Cycle 1 (C1). A total of 98 lines were selected from C1 based on genomic estimated breeding value (GEBV) and randomly intercrossed to generate the next cycle of progeny. Training populations used for genomic prediction and approaches for updating those populations are detailed in [41]. The progeny from the intercrosses among selected lines were selfed to the F_3_ generation. The process of line selection, intercrossing, and inbreeding, was repeated, creating three cycles of selection using genomic prediction. The total number of lines selected for C2 was 105, and the total number of lines selected for C3 was 48 (S1 Fig). Breeding program progress was evaluated by phenotypic comparison of the selected lines to a random subset of lines from each cycle. The numbers of randomly selected lines were 300, 101, 49 from C1, C2, and C3, respectively (S1 Fig).

Selection was based on the predicted phenotypic values for grain yield and for reduced fungal disease severity using a proxy phenotype, the concentration of the mycotoxin, deoxynivalenol (DON) which is created during an active Fusarium infection [41]. GEBV prediction was based on 384 SNPs evenly distributed across the seven barley chromosomes and chosen to maximize marker informativeness among the founders. Genotyping used an Illumina Veracode assay [41]. Lines were selected for increased yield and reduced DON concentration. GEBVs were estimated with ridge regression, as implemented in the ‘rrBLUP’ package [54] for R [55].

### Phenotypic Data Collection

F_3:5_ breeding lines in the selected and random pools for Cycles 1-3 were evaluated in yield trials at five year-locations. Phenotypic data was collected on grain yield and DON concentration. Phenotypic data were spatially adjusted with a moving average across the field plots. Best linear unbiased estimates (BLUEs) for yield and DON concentration were then produced for each line using the ‘rrBLUP’ package for R.

Raw and adjusted phenotypic data, including planting locations in the field trials, are available at https://github.com/MorrellLAB/Deleterious_GP and http://conservancy.umn.edu/XXXX. For details of phenotypic data collection see [see 41].

### Genotypic Data Collection

A total of 5,215 F_3_ progeny were genotyped across the three cycles using the 384 SNPs from the barley oligo pooled assay (BOPA) marker panel [56]. The physical location of all SNPs were determined based on automated BLAST searches against the barley reference genome [43], using consensus genetic map position to resolve ambiguous positions [57]. A small number of SNPs were missing either a genetic or physical position. For these SNPs we use linear interpolation as described in the [S2 Appendix].

Genotypes were called using signal to noise ratios from the raw probe intensities, as implemented in machine-scoring algorithm ALCHEMY [58]. ALCHEMY was used for genotype calls because it does not rely on clustering of samples to identify genotypic classes, thus avoiding Hardy-Weinberg equilibrium genotype frequency assumptions, and makes use of a prior estimate of the inbreeding coefficient to model the number of expected heterozygous genotypes. The prior inbreeding coefficient was specified as 0.99 for parental lines and as 0.75 for the F_3_ progeny, the average expected inbreeding coefficient after two generations of self-fertilization. Genotyping data was transformed to PLINK 1.9 format [59], and included pedigree information for each individual (data available at http://conservancy.umn.edu/XXXX). PLINK was used to test for Mendelian errors in inheritance of SNPs and to orient SNPs on the appropriate strand relative to the barley reference genome sequence from the cultivar Morex [43]. SNP genotypes from the barley BOPA markers genotyped in the Morex X Steptoe genetic mapping population [60,61] were used to infer the reference strand of origin for each SNP. The “hybrid peeling” approach of [45] was used for simultaneous imputation and phasing of genotyping data and thus to infer the parental contribution of chromosomal segments to progeny.

The approach of [45] makes use of an extended pedigree so that phased genotype inference is improved by comparisons to both progenitors and progeny. The specified pedigree is available at https://github.com/MorrellLAB/Deleterious_GP/blob/master/Data/Pedigrees/AlphaPeel_Pedigree.txt. PLINK was used for a second round of Mendel error checking with imputed genotypes. Imputed genotypes in progeny were set to missing if their genotype probability was less than 0.7.

### DNA Extraction, Sequence Analysis, and Variant Calling

DNA was extracted from young leaf tissue from each of the 21 founder lines using the Plant DNAzol extraction reagent and protocol from Thermo Fisher Scientific (Waltham, MA). Genomic DNA was captured with liquid phase exome probes designed to capture 60 Mb of the barley genome [62]. Eighteen of the samples were sequenced with 100 bp paired end technology on an Illumina HiSeq2000, and three were sequenced with 125 bp paired end technology on an Illumina HiSeq2500. Exomes were sequenced to a target depth of 30-fold coverage. Raw FASTQ files were cleaned of 3’ sequencing adapter contamination with Scythe (https://github.com/vsbuffalo/scythe), using a prior on contamination rate of 0.05. Adapter trimmed reads were then aligned to the Morex pseudo-molecule assembly (http://webblast.ipk-gatersleben.de/registration/) with BWA-MEM [63]. Mismatch and alignment reporting parameters were tuned to allow for approximately three high-quality mismatches between the reads and the reference. This represents approximately the highest observed coding sequence diversity in barley [64,65]. BAM files were cleaned of unmapped reads, split alignments, and sorted with SAMtools version 1.3 [66]. Duplicate reads were removed with Picard version 2.0.1 (http://broadinstitute.github.io/picard/).

Alignment processing followed the Genome Analysis Toolkit (GATK) best practices workflow [67,68]. Cleaned BAM alignments were realigned around putative insertion/deletion (indel) sites. Individual sample genotype likelihoods were then calculated with the HaplotypeCaller, with a haploid model and “heterozygosity” value of 0.008 per base pair. This value is the mean estimate of coding nucleotide sequence diversity, based on previous Sanger resequencing experiments [65,69]. SNP calls were made from the genotype likelihoods with the GATK tool GenotypeGVCFs [68].

Estimates of read depth and coverage made use of ‘bedtools genomecov’ relative to an empirical estimate of exome coverage. Briefly, estimated exome coverage was based on BWA-MEM mapping of roughly 241-fold exome capture reads from the reference barley line Morex (SRA accession number ERR271711), against the Morex draft genome. Read mapping was performed using the same parameters as for mapping the reads from the parental varieties against the reference assembly. Regions covered by at least 50 reads were considered covered by exome capture. Intervals that were separated by 50 bp or fewer were joined into a single interval. This results in ∼80 Mb of exome coverage relative to the 60 Mb based on capture probe design [62]. Recombination rate in cM/Mb was estimated based on physical positions of SNPs in the reference genome [43] and the estimated crossover rate from the consensus genetic map of [61].

Scripts to perform adapter contamination removal, read mapping, alignment cleaning, and implementing the GATK best practices workflow are available at https://github.com/MorrellLAB/Deleterious_GP. The BED file describing the empirical estimate of capture coverage is also available at the provided GitHub link and at http://conservancy.umn.edu/XXXX.

### Inference of Ancestral State Using an Outgroup Sequence

Whole genome resequencing data for *Hordeum murinum* ssp. *glaucum* was collected using Illumina paired end 150 bp reads on a NextSeq system. We chose *H. murinum* ssp. *glaucum* for ancestral state inference because phylogenetic analyses have placed this diploid species in a clade relatively close to *H. vulgare* [70]. Previous comparison of Sanger and exome capture resequencing from the most closely related species, *H. bulbosum*, identified shared polymorphisms at a proportion of SNPs, resulting in ambiguous ancestral states [65,65]. After adapter trimming, sequencing reads from *H. murinum* ssp. *glaucum* were mapped to the Morex reference genome using Stampy version 1.0.31 [44], with prior divergence estimates of 3%, 5%, 7.5%, 9%, and 11%. Cleaned BAM files were generated using Samtools version 1.3.1 [66] and Picard version 2.1.1 (http://broadinstitute.github.io/picard) and realigned around insertion/deletions (indel) using GATK version 3.6. A *H. murinum ssp. glaucum* FASTA file was created using ANGSD/ANGSD-wrapper [71,72]. Inference of ancestral state for SNPs in this set of 21 parents was performed using a custom Python script. For the above sequence processing pipeline, the following steps were performed using ‘sequence_handling’ [73] for quality control, adapter trimming, cleaning BAM files, and coverage summary. All other steps for processing *H. murinum ssp. glaucum* and inferring ancestral state are available on GitHub (https://github.com/liux1299/Barley_Outgroups).

### Deleterious Predictions

Variant annotation, including the identification of nonsynonymous variants used gene models provided by [43]. Annotations were applied to the reference genome using ANNOVAR [74]. Nonsynonymous SNPs were tested with three prediction approaches: PROVEAN [75], Polymorphism Phenotyping 2 (PPH2) [76], and BAD_Mutations [25,77] which implements a likelihood ratio test for neutrality [22]. All three approaches use phylogenetic sequence constraint to predict whether a base substitution is likely to be deleterious. PROVEAN and PPH2 used BLAST searches against the NCBI non-redundant protein sequence database, current as of 30 August, 2016. BAD_Mutations was run with a set of 42 publicly available Angiosperm genome sequences, hosted on Phytozome (https://phytozome.jgi.doe.gov) and Ensembl Plants (http://plants.ensembl.org/). A SNP was considered deleterious by PROVEAN if the substitution score was less than or equal to −4.1528, as determined by calculating 95% specificity from a set of known phenotype-altering SNPs in *Arabidopsis thaliana* [77]. PPH2 classifies SNPs as neutral or deleterious; prediction was considered as deleterious if it output a ‘deleterious’ call for a SNP. These programs use a heuristic for testing evolutionary constraint, as well as a training model for known human disease-causing polymorphisms. A SNP was considered deleterious by BAD_Mutations if the *p*-value from a logistic regression [24] was less than 0.05. The logistic regression model used for dSNP identification is an update to the BAD_Mutations implementation reported by [24]. For comparative analyses, nonsynonymous SNPs were considered to be deleterious if they were identified as deleterious by all three approaches, or if they form an early stop codon (nonsense SNP).

### Population Summary Statistics

Pairwise diversity across classes of SNPs was calculated using VCFTools and diploid genotypes for each individual. Calculations were partitioned across breeding cycles and among functional classes including noncoding, synonymous, nonsynonymous, and deleterious. For progeny in C1-C3, calculations made use of imputed genotypes relative to the transmission of the 384 SNP genotyping for each partition and functional class.

For SNP genotyping, the expected number of segregating markers in each family was calculated based on the pedigree and based on SNPs that were polymorphic within families based on parental genotypes.

Testing for segregation distortion at individual SNPs used the parental genotypes and the pedigrees to generate the expected number of genotypes. The observed numbers of each genotypic class in each cycle were calculated with PLINK. Observed genotype counts were tested for significant departure from Mendelian expectations using Fisher’s Exact Test, implemented in the R statistical computing environment [55].

### Proportion of Phenotypic Variance Explained

The proportion of phenotypic variance that could be explained from the genotyping data was estimated using linear mixed model approach as implemented in the program Genome-wide Efficient Mixed Model Association (GEMMA) [46]. The model incorporates estimated a identity matrix among samples that controls for family structure. We estimated the phenotypic variance explained for three phenotypes, yield, DON concentration, and plant height, using spatially adjusted BLUP (or BLUE) estimates averaged across years and locations as reported by [41].

The mixed model analysis involved a minimum minor allele frequency of 1% in the phenotyping panel. Heritability was estimated for SNPs from the 384 SNP Veracode panel and for SNPs imputed from parents to progeny based on exome capture resequencing.

## Acknowledgements

The authors are also grateful to Ana Poets and Tyler Tiede for sharing computer code used in data analysis. We are grateful to Maria Muñoz-Amatriaían and Timothy Close for providing SNP genotyping data from barley genetic mapping populations that included the variety Morex, used for the reference genome. We are grateful to Shiaoman Chao for provided barley genotyping raw data. Sebastian Beier provided physical positions for a portion of genotyped SNPs. Li Lei, Yong Jiang, Jochen Reif, Albert Schulthess, Ruth Shaw, Robert Stupar, Peter Tiffin, and Yusheng Zhao provided valuable comments on an earlier version of the text. This research was carried out with hardware and software support provided by the Minnesota Supercomputing Institute (MSI) at the University of Minnesota.

## Data Availability Statement

All raw sequence reads for barley parental lines are available as BioProject ID PRJNA399170. Raw reads for *Hordeum murinum* ssp. *glaucum* are available as BioProject ID PRJNA491526. Additional files including a variant call format (VCF) file of variants called in all parents, and empirical estimate of exome capture coverage, and all available genotypes and phenotypes are in a Data Repository University of Minnesota (DRUM) archive at http://conservancy.umn.edu/XXXX. Scripts used for analysis are available at https://github.com/MorrellLAB/Deleterious_GP. All other relevant data is found in the paper and its Supporting Information files.

## Author Contributions

**Conceptualization**: Thomas J. Y. Kono, Justin C. Fay, Kevin P. Smith, Peter L. Morrell

**Data curation**: Thomas J. Y. Kono, Emily E. Vonderharr, Daniel Koenig, Kevin P. Smith, Peter L. Morrell

**Formal analysis:** Thomas J. Y. Kono, Chaochih Liu, Peter L. Morrell

**Funding acquisition:** Justin C. Fay, Kevin P. Smith, Peter L. Morrell

**Investigation:** Thomas J. Y. Kono, Kevin P. Smith, Peter L. Morrell

**Methodology:** Thomas J. Y. Kono, Justin C. Fay, Peter L. Morrell

**Project administration:** Justin C. Fay, Kevin P. Smith, Peter L. Morrell

**Resources:** Daniel Koenig, Kevin P. Smith, Peter L. Morrell

**Software:** Thomas J. Y. Kono, Chaochih Liu, Justin C. Fay, Peter L. Morrell

**Supervision:** Justin C. Fay, Peter L. Morrell

**Validation:** Thomas J. Y. Kono, Chaochih Liu, Emily E. Vonderharr, Peter L. Morrell

**Visualization:** Thomas J. Y. Kono, Peter L. Morrell

**Writing – original draft:** Thomas J. Y. Kono, Chaochih Liu, Justin C. Fay, Peter L. Morrell

**Writing – review & editing:** Thomas J. Y. Kono, Chaochih Liu, Emily E. Vonderharr, Daniel Koenig, Justin C. Fay, Kevin P. Smith, Peter L. Morrell

---

S1 Fig. A schematic of the barley genomic prediction population used in this study.

S2 Fig. The folded site frequency spectra (SFS) for all SNPs in the parental founders and three cycles of progeny in the genomic prediction population. The SFS for progeny is imputed relative to genotyped SNPs. SNPs are partitioned by functional classes.

S3 Fig. Exome capture target density (dark blue line), recombination rate in cM/Mb (green line), and the genomic distribution of SNPs identified in the parental varieties (vertical light blue lines). Purple triangles indicate SNPs genotyped in parents and progeny. Exome capture target density is the number of exome capture targets per 100kb. Recombination rate estimates are derived from the genetic map of [61], and Lowess-smoothed in windows of 3Mb, using 2% of the points in each window for smoothing.

S4 Fig. The derived site frequency spectrum (SFS) for all sites with estimated ancestral state based on imputed genotypes relative to genotyped SNPs in progeny for each cycle. SNPs are partitioned by functional classes.

S5 Fig. The folded site frequency spectrum (SFS) for all imputed variants in all cycles. SNPs are partitioned by functional classes.

S6 Fig. Proportion of variants that fixed for ancestral states in C3 at various initial frequencies in the founders. Variants are partitioned by functional class.

S7 Fig. Fold change in derived allele frequency for all variants with unambiguous ancestral states from C1 to C3 across the genome. Grey shading indicates pericentromeric regions.

S8 Fig. Average burden of dSNPs carried by each individual in the population, measured as number of derived alleles at all identified deleterious sites.

## Supplemental Materials

## S1 Appendix Yield Trials

For yield trials in 2014, lines were evaluated at Crookston, MN; Morris, MN; and Saint Paul, MN. For 2015 yield trials, lines were evaluated at Crookston and Morris. Lines were grown in an augmented block design [78]. The check varieties were to adjust for spatial variation across trial plots. Checks included ‘Lacey’ (96 replicates), ‘Quest’ (24 replicates), ‘Stellar-ND’ (20 replicates), and ‘Tradition’ (20 replicates).

For DON concentration trials, each chosen F_3:5_ line was evaluated at five year-locations in disease nurseries [79]. Similar to the yield trials, lines were grown in an augmented block design. DON concentration was evaluated at Crookston, MN in 2013, 2014, and 2015. DON concentration was evaluated at Saint Paul, MN in 2013 and 2014. Check varieties for DON trials were ‘Quest’ (123 replicates), ‘ND20448’ (26 replicates), ‘Tradition’ (25 replicates), and ‘Lacey’ (25 replicates).

## S2 Appendix Linear Interpolation of Genetic and Physical SNP Positions

Among the 384 SNPs on the Veracode assay, two were missing genetic positions and 14 were missing physical positions. To interpolate genetic or physical positions, we use the positions of flanking SNPs. We take half the distance between known positions. In the formulas, D is average distance, G is genetic distance, P is physical distance, and the subscripts k and u refer to known and unknown positions and subscripts 1 and 2 refer to positions up and downstream of the position to be interpolated.

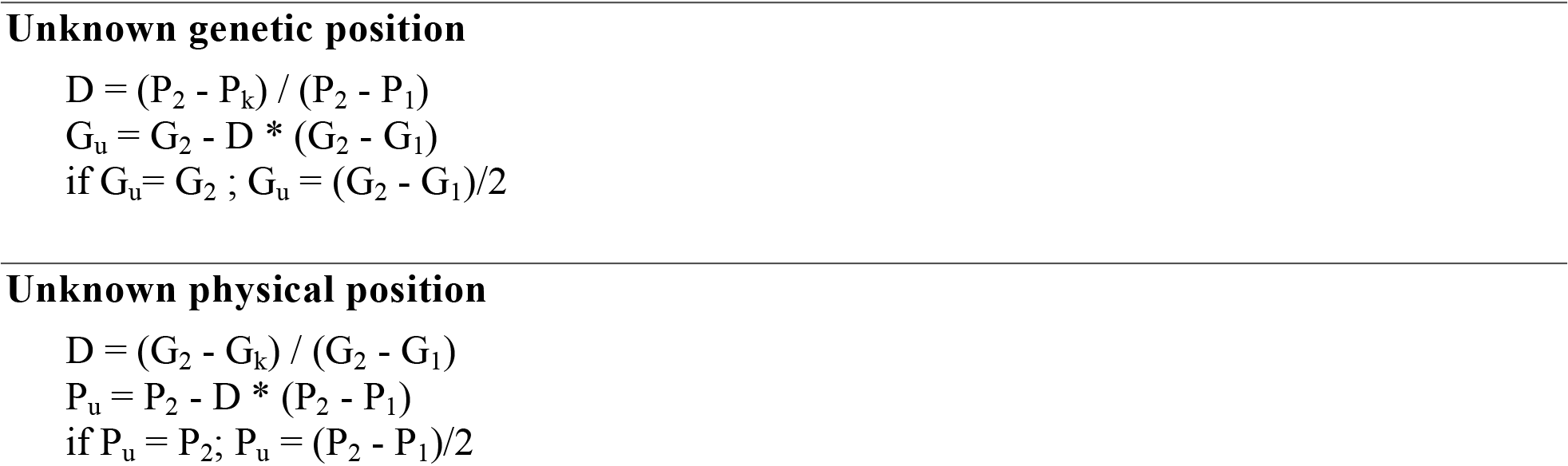

---

**S1 Table.**
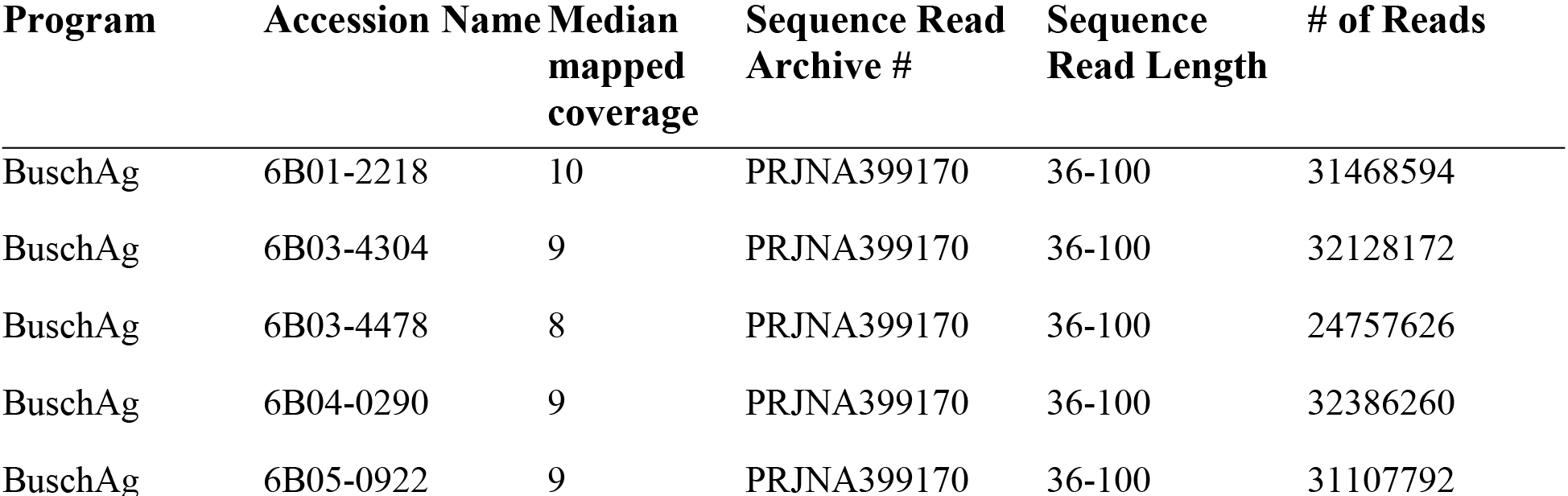
Founder parents used in the genomic prediction experiment along with exome capture resequencing summaries.

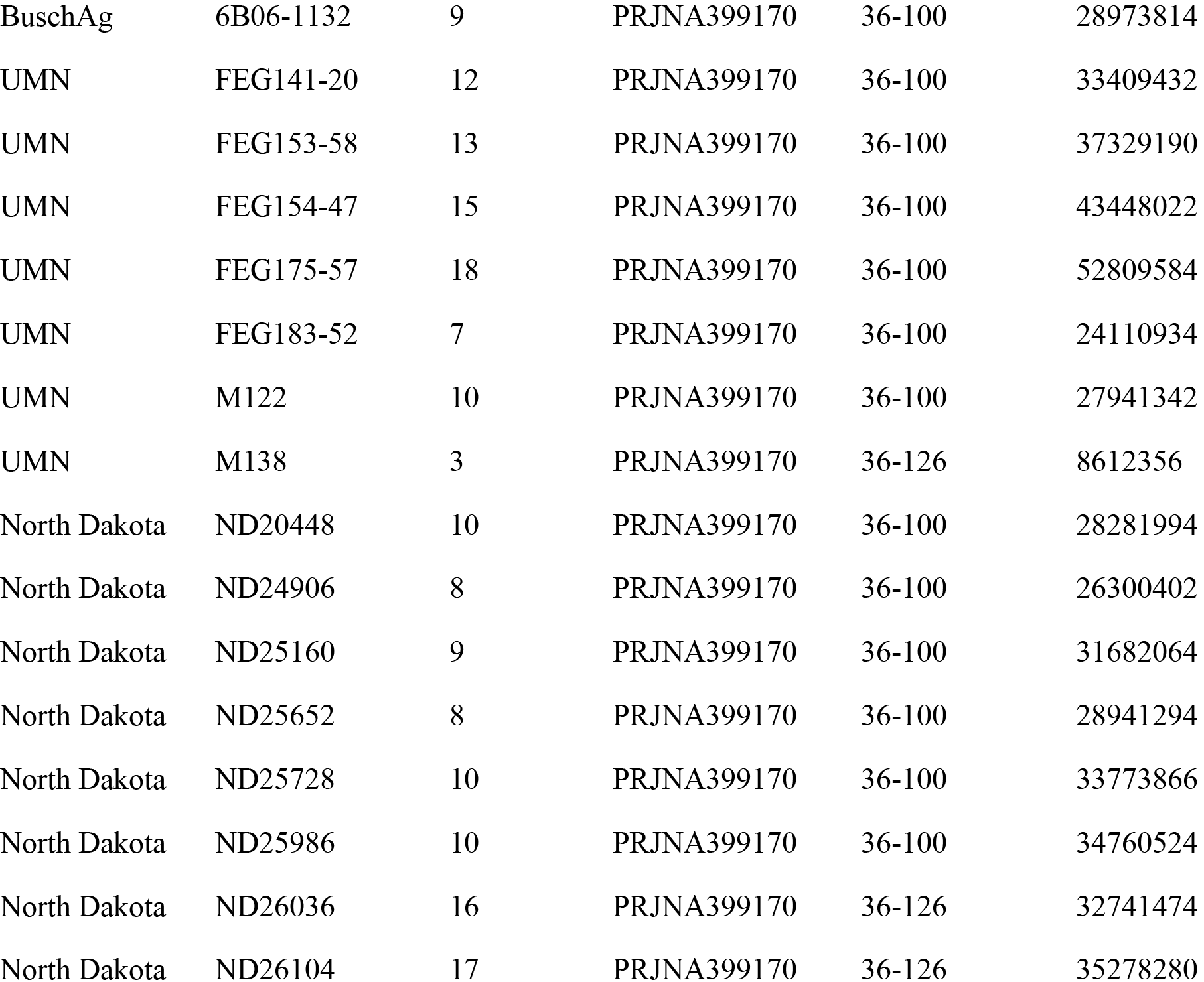

**S2 Table.** Summary of putatively dSNPs in the founder lines. BA: Busch Agricultural Resources, Inc.; MN: University of Minnesota; ND: North Dakota State University. Values listed include the number of dSNPs and private dSNPs per inbred line.

| Breeding Program | Line ID | dSNPs | Private dSNPs |
| --- | --- | --- | --- |
| BA | 6B01-2218 | 680 | 83 |
| BA | 6B03-4304 | 668 | 24 |
| BA | 6B03-4478 | 670 | 37 |
| BA | 6B04-0290 | 665 | 27 |
| BA | 6B05-0922 | 693 | 47 |
| BA | 6B06-1132 | 674 | 25 |
| MN | FEG141-20 | 690 | 107 |
| MN | FEG153-58 | 698 | 13 |
| MN | FEG154-47 | 673 | 18 |
| MN | FEG175-57 | 690 | 36 |
| MN | FEG183-52 | 695 | 38 |
| MN | M138 | 659 | 11 |
| MN | M122 | 686 | 15 |
| ND | ND20448 | 665 | 46 |
| ND | ND24906 | 664 | 40 |
| ND | ND25160 | 669 | 135 |
| ND | ND25652 | 633 | 32 |
| ND | ND25728 | 705 | 72 |
| ND | ND25986 | 690 | 23 |
| ND | ND26036 | 676 | 90 |
| ND | ND26104 | 688 | 172 |

**S3 Table.**
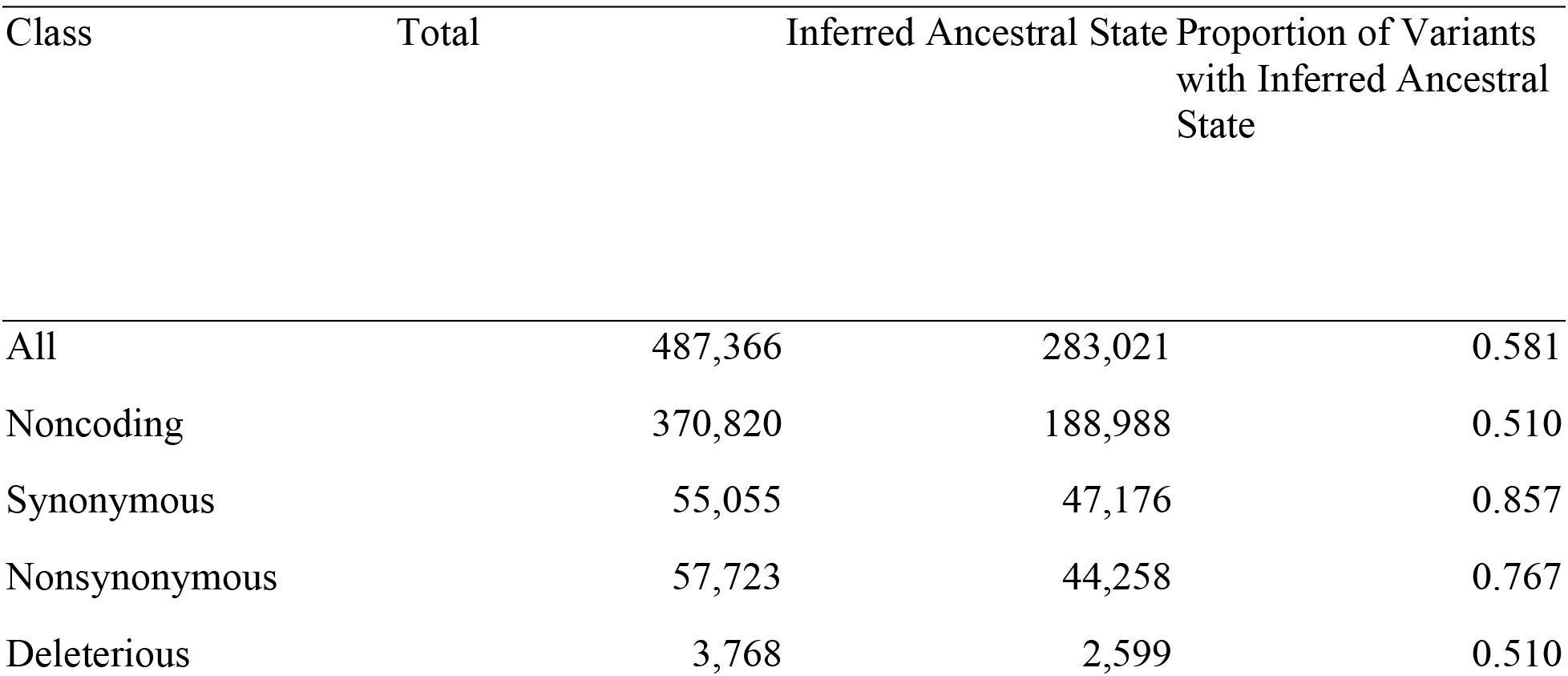
The ancestral state of all variants was inferred with *Hordeum murinum* ssp. *glaucum* used as an outgroup. The number of SNPs in each class and the proportion of SNPs for which ancestral state could be inferred is shown.

**S4 Table.**
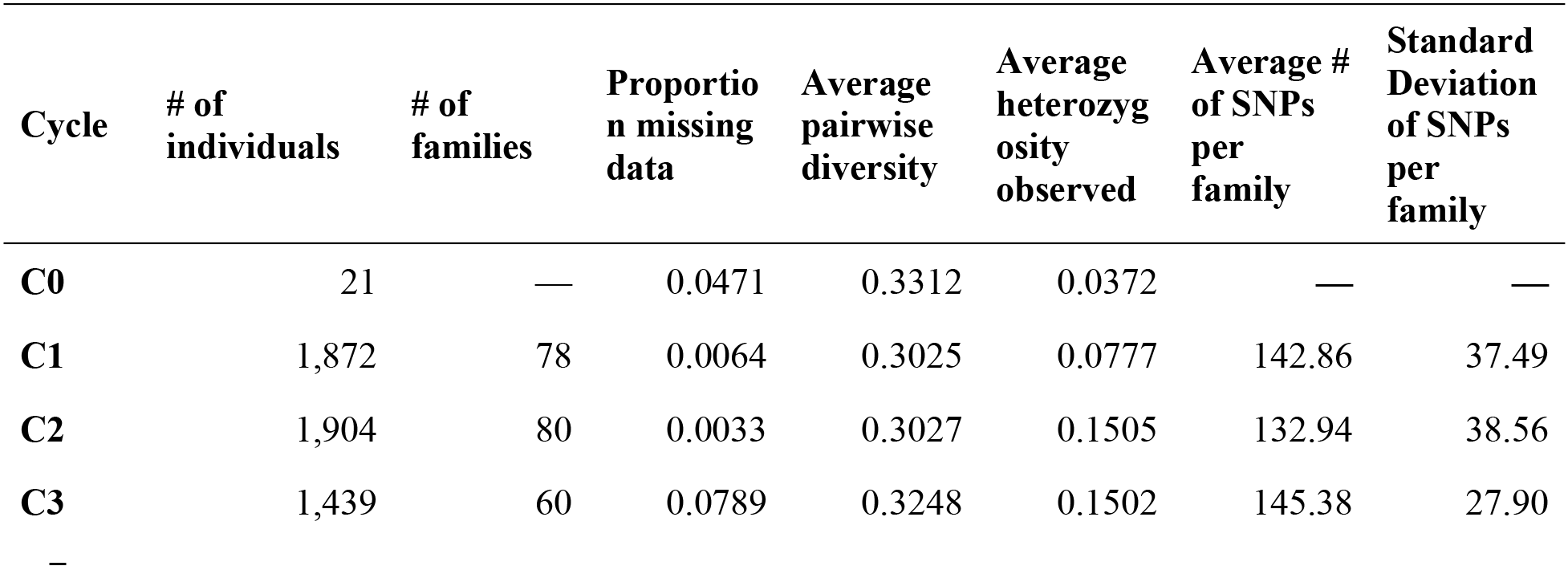
Basic descriptive statistics from genotyping of the Veracode 384 SNP assay. Values reported are based on observed genotypes in Illumina genotyping or exome capture resequencing (for C0 founder lines).

**S5 Table.** Average pairwise diversity among progeny for sites that were polymorphic in the C0 founder lines. The data is based on phased and imputed genotypes given observed genotypes from the Veracode 384 SNP assay.

| Cycle | All Sites | Noncoding | Synonymous | Nonsynonymous | Deleterious |
| --- | --- | --- | --- | --- | --- |
| All | 0.170 | 0.168 | 0.186 | 0.165 | 0.122 |
| C0 | 0.191 | 0.188 | 0.210 | 0.189 | 0.148 |
| C1 | 0.186 | 0.184 | 0.204 | 0.183 | 0.138 |
| C2 | 0.161 | 0.160 | 0.177 | 0.157 | 0.115 |
| C3 | 0.152 | 0.151 | 0.166 | 0.146 | 0.105 |

## References

1. Ewens WJ (1972) The sampling theory of selectively neutral alleles. Theor Popul Biol 3: 87–112.

2. Watterson GA (1975) On the number of segregating sites in genetical models without recombination. Theor Popul Biol 7: 256–276.

3. Thornton KR, Foran AJ, Long AD (2013) Properties and modeling of GWAS when complex disease risk is due to non-complementing, deleterious mutations in genes of large effect. PLoS Genet 9: e1003258.

4. Eyre-Walker A, Woolfit M, Phelps T (2006) The distribution of fitness effects of new deleterious amino acid mutations in humans. Genetics 173: 891–900.

5. Sanjuán R, Moya A, Elena SF (2004) The distribution of fitness effects caused by single-nucleotide substitutions in an RNA virus. Proc Natl Acad Sci USA 101: 8396–8401.

6. Thatcher JW, Shaw JM, Dickinson WJ (1998) Marginal fitness contributions of nonessential genes in yeast. Proceedings of the National Academy of Sciences 95: 253–257.

7. Johnson T, Barton N (2005) Theoretical models of selection and mutation on quantitative traits. Philos Trans R Soc Lond B Biol Sci 360: 1411–1425.

8. Eyre-Walker A (2010) Evolution in health and medicine Sackler colloquium: Genetic architecture of a complex trait and its implications for fitness and genome-wide association studies. Proc Natl Acad Sci USA 107 Suppl 1: 1752–1756.

9. Stanton-Geddes J, Paape T, Epstein B, Briskine R, Yoder J, Mudge J, Bharti AK, Farmer AD, Zhou P, Denny R, May GD, Erlandson S, Yakub M, Sugawara M, Sadowsky MJ, Young ND, Tiffin P (2013) Candidate genes and genetic architecture of symbiotic and agronomic traits revealed by whole-genome, sequence-based association genetics in Medicago truncatula. PLoS One 8: e65688.

10. Lewontin RC (1974) The genetic basis of evolutionary change. New York: Columbia University Press. xiii, 346 p.

11. Beavis WD (1994) The power and deceit of QTL experiments: lessons from comparative QTL studies. Proceedings of the forty-ninth annual corn and sorghum industry research conference: 250–266.

12. Rockman MV (2012) The QTN program and the alleles that matter for evolution: all that’s gold does not glitter. Evolution 66: 1–17.

13. Felsenstein J (1974) The evolutionary advantage of recombination. Genetics 78: 737–756.

14. Marth GT, Yu F, Indap AR, Garimella K, Gravel S, Leong WF, Tyler-Smith C, Bainbridge M, Blackwell T, Zheng-Bradley X, Chen Y, Challis D, Clarke L, Ball EV, Cibulskis K, Cooper DN, Fulton B, Hartl C, Koboldt D, Muzny D, Smith R, Sougnez C, Stewart C, Ward A, Yu J, Xue Y, Altshuler D, Bustamante CD, Clark AG, Daly M, DePristo M, Flicek P, Gabriel S, Mardis E, Palotie A, Gibbs R, 1000 GP (2011) The functional spectrum of low-frequency coding variation. Genome Biol 12: R84.

15. Lu J, Tang T, Tang H, Huang J, Shi S, Wu CI (2006) The accumulation of deleterious mutations in rice genomes: a hypothesis on the cost of domestication. Trends Genet 22: 126–131.

16. Ramu P, Esuma W, Kawuki R, Rabbi IY, Egesi C, Bredeson JV, Bart RS, Verma J, Buckler ES, Lu F (2017) Cassava haplotype map highlights fixation of deleterious mutations during clonal propagation. Nat Genet 49: 959–963.

17. Marsden CD, Ortega-Del Vecchyo D, O’Brien DP, Taylor JF, Ramirez O, Vilà C, Marques-Bonet T, Schnabel RD, Wayne RK, Lohmueller KE (2016) Bottlenecks and selective sweeps during domestication have increased deleterious genetic variation in dogs. Proceedings of the National Academy of Sciences 113: 152–157.

18. Zhou Y, Massonnet M, Sanjak JS, Cantu D, Gaut BS (2017) Evolutionary genomics of grape (Vitis vinifera ssp. vinifera) domestication. Proc Natl Acad Sci USA 114: 11715–11720.

19. Liu Q, Zhou Y, Morrell PL, Gaut BS (2017) Deleterious variants in Asian rice and the potential cost of domestication. Mol Biol Evol 34: 908–924.

20. Moyers BT, Morrell PL, McKay JK (2017) Genetic costs of domestication and improvement. J Hered 109: 103–116.

21. Makino T, Rubin CJ, Carneiro M, Axelsson E, Andersson L, Webster MT (2018) Elevated proportions of deleterious genetic variation in domestic animals and plants. Genome Biol Evol 10: 276–290.

22. Chun S, Fay JC (2009) Identification of deleterious mutations within three human genomes. Genome Res 19: 1553–1561.

23. Ng PC (2003) SIFT: predicting amino acid changes that affect protein function. Nucleic Acids Res 31: 3812–3814.

24. Kono TJY, Lei L, Shih CH, Hoffman PJ, Morrell PL (2017) Comparative genomics approaches accurately predict deleterious variants in plants. bioRxiv

25. Kono TJ, Fu F, Mohammadi M, Hoffman PJ, Liu C, Stupar RM, Smith KP, Tiffin P, Fay JC, Morrell PL (2016) The role of deleterious substitutions in crop genomes. Mol Biol Evol 33: 2307–2317.

26. Renaut S, Rieseberg LH (2015) The accumulation of deleterious mutations as a consequence of domestication and improvement in sunflowers and other Compositae crops. Mol Biol Evol 32: 2273–2283.

27. Morrell PL, Buckler ES, Ross-Ibarra J (2012) Crop genomics: advances and applications. Nat Rev Genet 13: 85–96.

28. Johnsson M, Gaynor RC, Jenko J, Goijanc G, de Koning D-J, Hickey JM (2018) Removal of alleles by genome editing — RAGE against the deleterious load.

29. Smith KP, Thomas W, Gutierrez L, Bull H (2018) Genomics-based barley breeding. In: Stein N, Muehlbauer GJ, editors. The Barley Genome.Cham, Switzerland: Springer. pp. 287–315.

30. Valluru R, Gazave EE, Fernandes SB, Ferguson JN, Lozano R, Hirannaiah P, Zuo T, Brown PJ, Leakey ADB, Gore MA, Buckler ES, Bandillo N (2018) Leveraging mutational burden for complex trait prediction in sorghum.

31. Mezmouk S, Ross-Ibarra J (2014) The pattern and distribution of deleterious mutations in maize. G3 4: 163–171.

32. Henn BM, Botigué LR, Peischl S, Dupanloup I, Lipatov M, Maples BK, Martin AR, Musharoff S, Cann H, Snyder MP, Excoffier L, Kidd JM, Bustamante CD (2015) Distance from sub-Saharan Africa predicts mutational load in diverse human genomes. Proceedings of the National Academy of Sciences

33. Henn BM, Botigué LR, Bustamante CD, Clark AG, Gravel S (2015) Estimating the mutation load in human genomes. Nat Rev Genet 16: 333–343.

34. Yang J, Mezmouk S, Baumgarten A, Buckler ES, Guill KE, McMullen MD, Mumm RH, Ross-Ibarra J (2017) Incomplete dominance of deleterious alleles contributes substantially to trait variation and heterosis in maize. PLoS Genet 13: e1007019.

35. Felsenstein J, Yokoyama S (1976) The evolutionary advantage of recombination. II. Individual selection for recombination. Genetics 83: 845–859.

36. Charlesworth B, Morgan MT, Charlesworth D (1993) The effect of deleterious mutations on neutral molecular variation. Genetics 134: 1289–1303.

37. Gazal S, Finucane HK, Furlotte NA, Loh PR, Palamara PF, Liu X, Schoech A, Bulik-Sullivan B, Neale BM, Gusev A, Price AL (2017) Linkage disequilibrium-dependent architecture of human complex traits shows action of negative selection. Nat Genet

38. Pardiñas AF, Holmans P, Pocklington AJ, Escott-Price V, Ripke S, Carrera N, Legge SE, Bishop S, Cameron D, Hamshere ML, Han J, Hubbard L, Lynham A, Mantripragada K, Rees E, MacCabe JH, McCarroll SA, Baune BT, Breen G, Byrne EM, Dannlowski U, Eley TC, Hayward C, Martin NG, McIntosh AM, Plomin R, Porteous DJ, Wray NR, Caballero A, Geschwind DH, Huckins LM, Ruderfer DM, Santiago E, Sklar P, Stahl EA, Won H, Agerbo E, Als TD, Andreassen OA, B$kvad-Hansen M, Mortensen PB, Pedersen CB, Børglum AD, Bybjerg-Grauholm J, Djurovic S, Durmishi N, Pedersen MG, Golimbet V, Grove J, Hougaard DM, Mattheisen M, Molden E, Mors O, Nordentoft M, Pejovic-Milovancevic M, Sigurdsson E, Silagadze T, Hansen CS, Stefansson K, Stefansson H, Steinberg S, Tosato S, Werge T, Gerad1 C, Crestar C, Collier DA, Rujescu D, Kirov G, Owen MJ, O’Donovan MC, Walters JTR, Gerad1 C, Crestar C, Gerad1 C, Crestar C (2018) Common schizophrenia alleles are enriched in mutation-intolerant genes and in regions under strong background selection. Nat Genet

39. Meuwissen THE, Hayes BJ, Goddard ME (2001) Prediction of total genetic value using genome-wide dense marker maps. Genetics 157: 1819–1829.

40. Edwards SM, Sørensen IF, Sarup P, Mackay TF, Sørensen P (2016) Genomic prediction for quantitative traits is improved by mapping variants to gene ontology categories in Drosophila melanogaster. Genetics 203: 1871–1883.

41. Tiede T, Smith KP (2018) Evaluation of retrospective optimization of genomic selection for yield and disease resistance in spring barley. Mol Breed

42. Muñoz-Amatriaín M, Lonardi S, Luo M, Madishetty K, Svensson JT, Moscou MJ, Wanamaker S, Jiang T, Kleinhofs A, Muehlbauer GJ, Wise RP, Stein N, Ma Y, Rodriguez E, Kudrna D, Bhat PR, Chao S, Condamine P, Heinen S, Resnik J, Wing R, Witt HN, Alpert M, Beccuti M, Bozdag S, Cordero F, Mirebrahim H, Ounit R Wu Y, You F, Zheng J, Simková H, Dolezel J, Grimwood J, Schmutz J, Duma D, Altschmied L, Blake T, Bregitzer P, Cooper L, Dilbirligi M, Falk A, Feiz L, Graner A, Gustafson P, Hayes PM, Lemaux P, Mammadov J, Close TJ (2015) Sequencing of 15 622 gene-bearing BACs clarifies the gene-dense regions of the barley genome. Plant J 84: 216–227.

43. Mascher M, Gundlach H, Himmelbach A, Beier S, Twardziok SO, Wicker T, Radchuk V, Dockter C, Hedley PE, Russell J, Bayer M, Ramsay L, Liu H, Haberer G, Zhang XQ, Zhang Q, Barrero RA, Li L, Taudien S, Groth M, Felder M, Hastie A, Šimková H, Staňková H, Vrána J, Chan S, Muňoz-Amatriain M, Ounit R, Wanamaker S, Bolser D, Colmsee C, Schmutzer T, Aliyeva-Schnorr L, Grasso S, Tanskanen J, Chailyan A, Sampath D, Heavens D, Clissold L, Cao S, Chapman B, Dai F, Han Y, Li H, Li X, Lin C, McCooke JK, Tan C, Wang P, Wang S, Yin S, Zhou G, Poland JA, Bellgard MI, Borisjuk L, Houben A, Doležel J, Ayling S, Lonardi S, Kersey P, Langridge P, Muehlbauer GJ, Clark MD, Caccamo M, Schulman AH, Mayer KFX, Platzer M, Close TJ, Scholz U, Hansson M, Zhang G, Braumann I, Spannagl M, Li C, Waugh R Stein N (2017) A chromosome conformation capture ordered sequence of the barley genome. Nature 544: 427–433.

44. Lunter G, Goodson M (2011) Stampy: a statistical algorithm for sensitive and fast mapping of Illumina sequence reads. Genome Res 21: 936–939.

45. Whalen A, Ros-Freixedes R Wilson DL, Gorjanc G, Hickey JM (2017) Hybrid peeling for fast and accurate calling, phasing, and imputation with sequence data of any coverage in pedigrees. bioRxiv

46. Zhou X, Stephens M (2012) Genome-wide efficient mixed-model analysis for association studies. Nat Genet 44: 821–824.

47. Fang Z, Eule-Nashoba A, Powers C, Kono TJY, Takuno S, Morrell PL, Smith KP (2013) Comparative analyses identify the contributions of exotic donors to disease resistance in a barley experimental population. G3: Genes, Genomes, Genetics 3: 1945–1953.

48. Tinker NA, Mather DE, Rossnagel BG, Kasha KJ, Kleinhofs A, Hayes PM, Falk DE, Ferguson T, Shugar LP, Legge WG (1996) Regions of the genome that affect agronomic performance in two-row barley. Crop Sci 36: 1053–1062.

49. Urrea CA, Horsley RD, Steffenson BJ, Schwarz PB (2002) Heritability of Fusarium head blight resistance and deoxynivalenol accumulation from barley accession CIho 4196. Crop Sci 42: 1404–1408.

50. Rodgers-Melnick E, Bradbury PJ, Elshire RJ, Glaubitz JC, Acharya CB, Mitchell SE, Li C, Li Y, Buckler ES (2015) Recombination in diverse maize is stable, predictable, and associated with genetic load. Proceedings of the National Academy of Sciences 112: 3823–3828.

51. Marais G, Charlesworth B, Wright SI (2004) Recombination and base composition: the case of the highly self-fertilizing plant Arabidopsis thaliana. Genome Biol 5: R45.

52. Brandvain Y, Wright SI (2016) The limits of natural selection in a nonequilibrium world. Trends Genet 32: 201–210.

53. Eyre-Walker A, Keightley PD (2007) The distribution of fitness effects of new mutations. Nat Rev Genet 8: 610–618.

54. Endelman JB (2011) Ridge regression and other kernels for genomic selection with R package rrBLUP. The Plant Genome Journal 4: 250.

55. Team RC (2018) R: A language and environment for statistical computing [Internet]. Vienna, Austria: R Foundation for Statistical Computing; 2018. Available: https://www.R-project.org/viatheInternet. Accessed x.

56. Close TJ, Bhat PR, Lonardi S, Wu Y, Rostoks N, Ramsay L, Druka A, Stein N, Svensson JT, Wanamaker S, Bozdag S, Roose ML, Moscou MJ, Chao S, Varshney RK, Szucs P, Sato K, Hayes PM, Matthews DE, Kleinhofs A, Muehlbauer GJ, DeYoung J, Marshall DF, Madishetty K, Fenton RD, Condamine P, Graner A, Waugh R (2009) Development and implementation of high-throughput SNP genotyping in barley. BMC Genomics 10: 582.

57. Lei L, Poets AM, Liu C, Wyant SR, Hoffman PJ, Carter CK, Trantow RM, Shaw BG, Li X, Muehlbauer G, Katagiri F, Morrell PL (2018) Discovery of barley gene candidates for low temperature and drought tolerance via environmental association.

58. Wright MH, Tung CW, Zhao K, Reynolds A, McCouch SR, Bustamante CD (2010) ALCHEMY: a reliable method for automated SNP genotype calling for small batch sizes and highly homozygous populations. Bioinformatics 26: 2952–2960.

59. Chang CC, Chow CC, Tellier LC, Vattikuti S, Purcell SM, Lee JJ (2015) Second-generation PLINK: rising to the challenge of larger and richer datasets. Gigascience 4: 7.

60. Kleinhofs A, Kilian A, Maroof MAS, Biyashev RM, Hayes P, Chen FQ, Lapitan N, Fenwick A, Blake TK, Kanazin V, Ananiev E, Dahleen L, Kudrna D, Bollinger J, Knapp SJ, Liu B, Sorrells M, Heun M, Franckowiak JD, Hoffman D, Skadsen R Steffenson BJ (1993) A molecular, isozyme and morphological map of the barley (Hordeum vulgare) genome. Theor Appl Genet 86: 705–712.

61. Muñoz-Amatriaín M, Moscou MJ, Bhat PR, Svensson JT, Bartoš J, Suchánková P, Šimková H, Endo TR Fenton RD, Lonardi S, Castillo AM, Chao S, Cistué L, Cuesta-Marcos A, Forrest KL, Hayden MJ, Hayes PM, Horsley RD, Makoto K, Moody D, Sato K, Vallés MP, Wulff BBH, Muehlbauer GJ, Doležel J, Close* TJ (2011) An improved consensus linkage map of barley based on flow-sorted chromosomes and single nucleotide polymorphism markers. The Plant Genome Journal 4: 238.

62. Mascher M, Richmond TA, Gerhardt DJ, Himmelbach A, Clissold L, Sampath D, Ayling S, Steuernagel B, Pfeifer M, D’Ascenzo M, Akhunov ED, Hedley PE, Gonzales AM, Morrell PL, Kilian B, Blattner FR Scholz U, Mayer KF, Flavell AJ, Muehlbauer GJ, Waugh R, Jeddeloh JA, Stein N (2013) Barley whole exome capture: a tool for genomic research in the genus Hordeum and beyond. Plant J 76: 494–505.

63. Li H, Durbin R (2009) Fast and accurate short read alignment with Burrows-Wheeler transform. Bioinformatics 25: 1754–1760.

64. Morrell PL, Toleno DM, Lundy KE, Clegg MT (2006) Estimating the contribution of mutation, recombination and gene conversion in the generation of haplotypic diversity. Genetics 173: 1705–1723.

65. Morrell PL, Gonzales AM, Meyer KK, Clegg MT (2014) Resequencing data indicate a modest effect of domestication on diversity in barley: a cultigen with multiple origins. J Hered 105: 253–264.

66. Li H, Handsaker B, Wysoker A, Fennell T, Ruan J, Homer N, Marth G, Abecasis G, Durbin R, 1000 GPDPS (2009) The Sequence Alignment/Map format and SAMtools. Bioinformatics 25: 2078–2079.

67. DePristo MA, Banks E, Poplin R, Garimella KV, Maguire JR, Hartl C, Philippakis AA, del Angel G, Rivas MA, Hanna M, McKenna A, Fennell TJ, Kernytsky AM, Sivachenko AY, Cibulskis K, Gabriel SB, Altshuler D, Daly MJ (2011) A framework for variation discovery and genotyping using next-generation DNA sequencing data. Nat Genet 43: 491–498.

68. McKenna A, Hanna M, Banks E, Sivachenko A, Cibulskis K, Kernytsky A, Garimella K, Altshuler D, Gabriel S, Daly M, DePristo MA (2010) The Genome Analysis Toolkit: a MapReduce framework for analyzing next-generation DNA sequencing data. Genome Res 20: 1297–1303.

69. Caldwell KS, Russell J, Langridge P, Powell W (2006) Extreme population-dependent linkage disequilibrium detected in an inbreeding plant species, Hordeum vulgare. Genetics 172: 557–567.

70. Jakob SS, Meister A, Blattner FR (2004) The considerable genome size variation of Hordeum species (Poaceae) is linked to phylogeny, life form, ecology, and speciation rates. Mol Biol Evol 21: 860–869.

71. Durvasula A, Kent TV, Hoffman PJ, Liu C, Kono TJY, Morrell PL, Ross-Ibarra J (2016) ANGSD-wrapper: utilities for analyzing next generation sequencing data. Molecular Ecology Resources 16: 1449–1454.

72. Korneliussen T, Albrechtsen A, Nielsen R (2014) ANGSD: Analysis of Next Generation Sequencing Data. BMC Bioinformatics 15: 356.

73. Hoffman PJ, Wyant SR, Kono TJY, Morrell PL (2018) MorrellLab/sequence_handling: Release v2.0: SNP calling with GATK 3.8. Available: https://zenodo.org/record/1257692#.WxLNny2ZOL5viatheInternet. Accessed x.

74. Wang K, Li M, Hakonarson H (2010) ANNOVAR: functional annotation of genetic variants from high-throughput sequencing data. Nucleic Acids Res 38: e164.

75. Choi Y, Sims GE, Murphy S, Miller JR, Chan AP (2012) Predicting the functional effect of amino acid substitutions and indels. PLoS One 7: e46688.

76. Adzhubei I, Jordan DM, Sunyaev SR (2013) Predicting functional effect of human missense mutations using PolyPhen-2. Curr Protoc Hum Genet Chapter 7: Unit7. 20.

77. Kono TJY, Lei L, Shih C-H, Hoffman PJ, Morrell PL, Fay JC (2018) Comparative genomics approaches accurately predict deleterious variants in plants. G3: Genes, Genomes, Genetics g3.200563.2018.

78. Lin C-S, Poushinsky GREG (1985) A modified augmented design (type 2) for rectangular plots. Can J Plant Sci 65: 743–749.

79. Massman J, Cooper B, Horsley R, Neate S, Dill-Macky R, Chao S, Dong Y, Schwarz P, Muehlbauer GJ, Smith KP (2011) Genome-wide association mapping of Fusarium head blight resistance in contemporary barley breeding germplasm. Mol Breeding 27: 439–454.

